# Damaging the conical morphology of HIV-1 capsid by targeting the FG-binding pocket and disfavoring pentameric subunits needed for core closure

**DOI:** 10.64898/2026.03.19.712970

**Authors:** William M. McFadden, Karen A. Kirby, Zachary C. Lorson, Lei Wang, Carolyn M. Highland, Sophie R. Harvey, Savannah Brancato, Andres Emanuelli Castaner, Haijuan Du, Vicki H. Wysocki, Zhengqiang Wang, Robert A. Dick, Stefan G. Sarafianos

## Abstract

The HIV-1 capsid is an essential viral component, targeted by the long-acting antiretroviral Lenacapavir (LEN). LEN binds to the HIV-1 capsid protein (CA) at the phenylalanine-glycine (FG) binding pocket (FGBP), a site for multiple host-factor and antiviral interactions in CA hexamers (CA_HEX_). Previously, we generated a chemical library to investigate the FGBP; ZW-1261, a lead compound, exhibits potent antiviral activity and strong inter-subunit interactions within CA_HEX_. Here, we report the molecular mechanism by which ZW-1261 affects the morphology and integrity of capsid lattice. ZW-1261 alone rapidly induces tubular CA assemblies; simultaneous addition of ZW-1261 with the assembly cofactor inositol hexaphosphate (IP6) forms morphologically distinct tubes. In mature virions, IP6 is required for the assembly of both CA_HEX_ and CA pentamers (CA_PENT_). Cryogenic-electron microscopy analysis of *in vitro* assembled capsid-like particles (CLPs) with IP6 suggests that ZW-1261 leads to the absence of CA_PENT_ and damages the pre-formed conical lattice. To elucidate how this FGBP-targeting antiviral impacts CA_PENT_, we further solved structures of CA_PENT_-only icosahedral assemblies (T = 1), formed by reported mutations, that were treated with ZW-1261. We find that ZW-1261 binding in these constrained T = 1 assemblies converts CA_PENT_ to a CA_HEX_-like conformation. Collectively, this suggests a mechanism by which addition of FGBP-binding inhibitor to native cores leads to the absence of CA_PENT_, impacting capsid closure and core integrity.

## Introduction

There are multiple classes of antiretroviral compounds that target various components of the human immunodominance virus type 1 (HIV-1) to inhibit viral replication (1, 2). Combinations of drugs within or between classes has been critical for the prevention of antiviral resistance for HIV-1, since resistance can lead to viral rebound and eventually the development of acquired immunodeficiency syndrome (AIDS) for people living with HIV (PLWH) taking antiretroviral therapies (ART), as well as increased potential for breakthrough infections for individuals at-risk of HIV-1 exposure taking protective antiretrovirals as pre-exposure prophylaxis (PrEP) (3–8). A drug with incredible potential to reshape the treatment and prevention of HIV-1 globally is the long-acting injectable Lenacapavir (LEN), a first-of-its-class antiretroviral approved for both ART and PrEP (1, 4, 9–12).

LEN is the first approved drug to target the HIV-1 capsid and is a highly potent inhibitor with a 50% effective concentration (EC_50_) ranging from 32–190 pM (1, 10, 11, 13–15). For HIV-1, the capsid is a great drug target due to its numerous, essential roles in viral replication, evident by its genetic fragility (4, 16, 17). The CA domain is one of the most well-conserved regions in the virus, as mutations that increase and that decrease the stability of the lattice have a negative impact on viral fitness; the capsid is a molecular container that encloses the viral genome, and as such, it is a protective shield that prevents innate immune sensing but consequently must also release the genome (“uncoating”) to enable integration at the correct time (17–25). Thus, compounds that modify the stability of the CA lattice have shown antiviral effects (10, 16, 26–29). The HIV-1 capsid also functions as vessel for reverse transcription and is a hub for host-cell factor interactions that facilitate immune evasion and intracellular trafficking, all of which are linked to the integrity and morphology of the core (23, 24, 27, 30–33). The morphology of the mature capsid is a determinant of its transport through the nuclear pore complex (NPC), a step critical for delivering the viral genome to the host chromatin for integration (22, 26, 30, 31, 34, 35). The mature capsid core is over 30 megadaltons in size and forms a fullerene-like cone shape, built of ∼1500 CA monomers that are arranged as ∼250 hexamers (CA_HEX_), which elongate the conical frustum, and exactly 12 pentamers (CA_PENT_), which provide curvature to the lattice and close the container (Figure 1A-1C) (27, 36–40). The CA monomers are the same sequence within CA_HEX_ and CA_PENT_ and are nearly identical structurally, with the major difference being at residues 58-TVGG-61 located at the base of α-helix 3 (α3) (41). For CA_HEX_, the monomers α3 span from Pro48 to Asn57 and have a disordered loop from 58-61, but for CA_PENT_, the monomers have elongated α3 to Gly60 (Figure 1D) (41–44). Mutating the two glycine residues to favor the elongated α3 with G60A/G61P enable the constitutive pentameric construct that form T = 1 icosahedrons (41).

**Figure 1:**
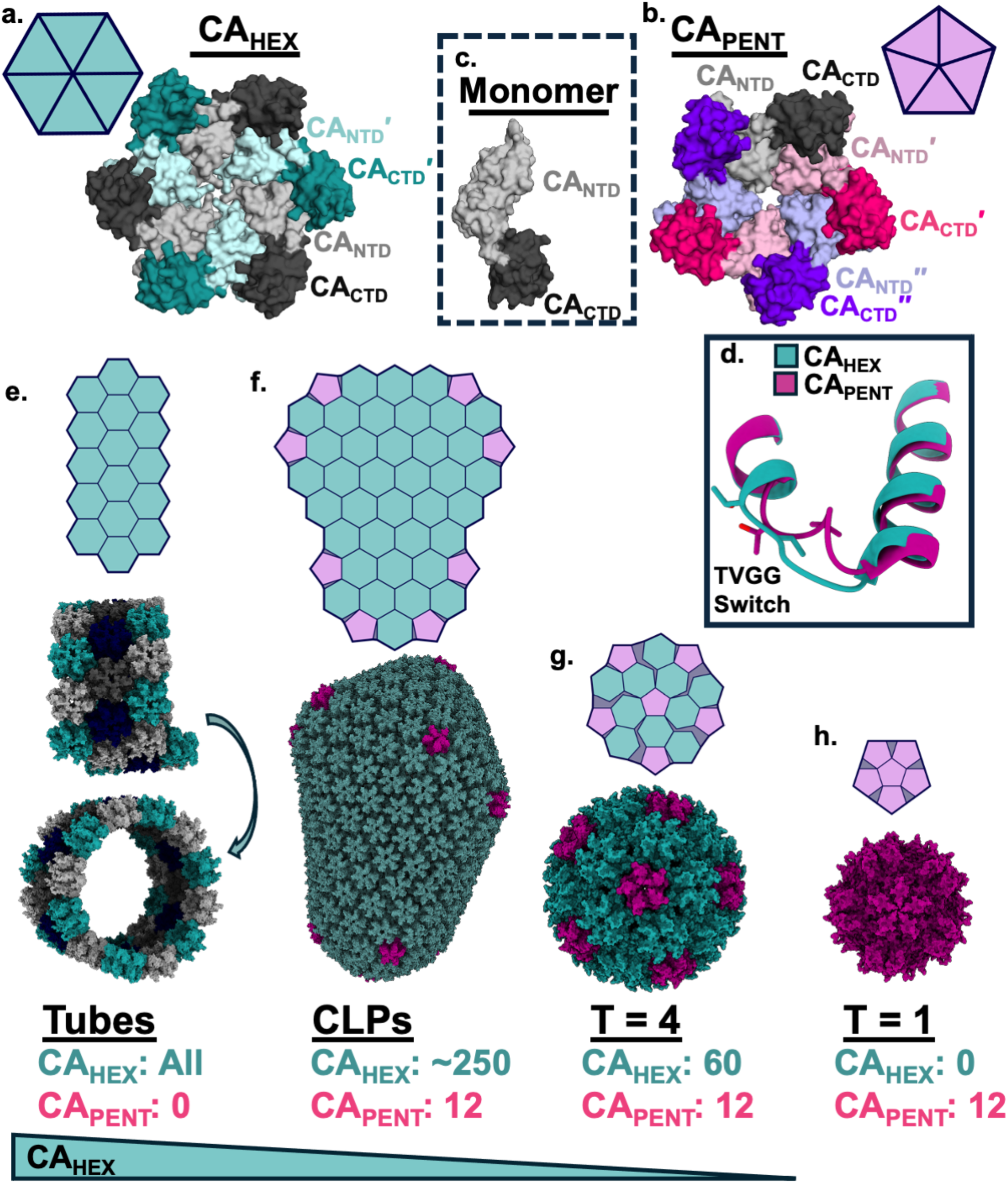
The geometric shape of macromolecular CA assemblies informs the composition of the CA_HEX_ and CA_PENT_ subunits. CA monomers can form **A.** hexameric (CA_HEX_) [PDB: 4XFX (42)] and/or **B.** pentameric (CA_PENT_) [PDB: 3P05 (87)] capsomers to build a macromolecular lattice from **C.** quasi-equivalent capsid protein (CA) monomers. **D.** The structural switch that is the structural difference in CA between CA_HEX_ (cyan) [PDB: 4XFX (42)] and CA_PENT_ (pink) [PDB: 8EEP (41)]. **E.** When CA_HEX_ (cyan, hexagons) exclusively interact, open-tubes or cylinders form that do not close [PDB: 3J4F (88)]. The addition of CA_PENT_ (pink, pentagons) adds curvature to the tubes and enables closure. **F.** For native fullerene-like cones to form as they do in the infectious virus, referred to as “capsid-like particles” (CLPs), CA assembles into approximately 250 CA_HEX_ with exactly 12 CA_PENT_ at the curved ends [PDB: 3J3Q (88)]. **G-H.** Mutations have been reported to decrease the prominence of CA_HEX_ in assemblies, such as **G.** N21C/A22C that can form T = 4 icosahedrons of 40 nm diameter with 60 CA_HEX_ and 12 CA_PENT_ [PDB: 4XFX, 3P05; EMDB-9733 (42, 50, 87)] or **H.** G60A/G61P that can form T = 1 icosahedrons of 20 nm diameter with no (0) CA_HEX_ and 12 CA_PENT_ [PDB: 8EEP (41)].

In the absence of CA_PENT_, a CA_HEX_-only assembly is tubular with open ends (Figure 1E) (35, 45, 46). Soluble assemblies with CA_PENT_ form an enclosed lattice due to the geometric curvature; in fullerene-like mature capsids or capsid-like particles, five CA_PENT_ are at the 40 nm narrow-end and seven CA_PENT_ are found at the 60 nm wide-end (Figure 1F) (47–49). Reducing the number of CA_HEX_ in the lattice through mutations enables icosahedral CA assemblies, where a 40 nm T = 4 icosahedron forms at a ratio of 60:12 CA_HEX_:CA_PENT_ and in the total absence of CA_HEX_ the 12 CA_PENT_ will form a 20 nm T = 1 icosahedron (Figure 1G-H) (38, 41, 50, 51).

The CA_HEX_ and CA_PENT_ multimers form distinct pockets that interact with different cellular proteins and cofactors, and these pockets have been targeted by antiviral compounds (Reviewed in: (27)). For transport into and through the NPC, the capsid interacts with host proteins that encode a phenylalanine-glycine (FG) motifs, like Nup153, Nup98, and CPSF6 that bind the CA_HEX_ FG binding pocket (FGBP) (19, 26, 52–55). The FGBP is a well-conserved hydrophobic pocket found between two monomers of a CA_HEX_, making six pockets for each multimer; it has been shown that unliganded CA_PENT_ have this pocket sterically blocked possibly preventing FG-interactions (27, 41, 42, 52). The FGBP is where LEN binds in mature cores, establishing this pocket as a clinically-relevant interface, and many other antivirals like PF74 and BI-2 bind here to modify capsid stability and can compete with host-factor binding (10, 14, 16, 26, 42, 52, 56–62). PF74 is a well-characterized small molecule inhibitor with three aromatic moieties (“R1”, “R2”, and “R3”), one of which mimics the FG motif of the host factors (“R2”; Supplementary Figure S1) (16, 19, 26, 27, 52, 59, 63). PF74 binds in-between CA N-terminal domain (CA_NTD_) and the C-terminal domain (CA_CTD_) with sub-µM affinity, but it has also been shown to interact with the CA_NTD_ alone with lower affinity and a different binding mode (26, 27, 42). Structure-based iterative drug design of the molecule PF74 has resulted in many FGBP-targeting antivirals, with ZW-1261 (Supplementary Figure S1) being a lead that exhibits potent antiviral effects (EC_50_ = 22 nM) and a favorable resistance profile (13, 27, 29, 60, 63, 64).

Another critical interface for CA_HEX_ and CA_PENT_ is Site 5 in the middle of the multimers at the central pore that interacts with acidic metabolites (Supplementary Figure S1), like inositol hexaphosphate (IP6), and some host-proteins with acidic domains, like PQBP1 (27, 61, 62, 65–69). This cationic channel is formed primarily by rings of basic residues, Arg18 and Lys25, that interact with the polyanionic chemicals like the dNTPs that fuel reverse transcription and IP6 molecules that enables mature core morphology (32, 44, 65–67, 70). IP6, or other polyanions, are required as cofactors for CA_PENT_ formation during capsid assembly, and thus, without acidic interactions at the central pore during maturation, there are only CA_HEX_ that cannot close the core (32, 49, 65–67, 70–72). While there is only one central pore for each multimer, it has been shown that multiple acidic chemicals can bind to this pocket simultaneously (32, 49, 70). Modifying this pocket can impact mature core formation and its stability, and clogging the pore preventing dNTP entry can decrease or impair reverse transcription (24, 32, 49, 70).

Here, we detail the structural relationship between the two neighboring pockets in capsid cores, the FGBP and the central pore, in both CA_HEX_ and CA_PENT_. We find that FGBP-targeting compounds favor CA_HEX_ and counter the established effects of polyanionic compounds that favor CA_PENT_. Native mass spectrometry (nMS) was used to determine the stoichiometry of FGBP and polyanionic ligands alone or in combination, and thermal shift assays (TSAs) helped elucidate the stabilizing nature of these combinations for CA_HEX_. ZW-1261 alone binds to and rapidly promotes CA_HEX_ assemblies, even in the absence of salt and at pH 8.0 that would typically prevent lattice formation. Further, ZW-1261 is a stronger inducer of assembly, more so than LEN or other PF74 analogs that we hypothesize is due to the modified indole ring of ZW-1261 (“R3”, Supplementary Figure S1) that can bridge two monomers of the CA_HEX_ by a coordinated water molecule (60).

Once a CA_PENT_ is formed, we find ZW-1261 can overcome steric blockage in FGBP open by the modifying an elongated α3 and create a CA_HEX_-like monomer that were arranged as CA_PENT_, but is dependent on the ionic strength of IP6 interactions. For soluble capsid-like particles (CLPs) assembled by excessive IP6 treatment (49), single-particle cryogenic electron microscopy (cryo-EM) found the compound bound to CA_HEX_ following but there were no CA_PENT_ classes after ZW-1261 treatment. Using the soluble CA_PENT_-only T = 1 icosahedrons enabled us to study the interactions of ZW-1261 in CA_PENT_ (41, 49). After forming the T = 1 icosahedron, addition of ZW-1261 would break the elongated α3 and revert the mutated molecular switch to that of a wild-type CA_HEX_. This aligns with reports about the mechanism of LEN impacting CA_PENT_ (73–75). Overall, we detail here the seemingly-antagonistic relationship between the polyanion IP6 that favor CA_PENT_ for capsid closure and the FGBP-targeting antiviral compounds that favor CA_HEX_ for lattice elongation, and how each ligand influences the morphology and integrity of the mature HIV-1 capsid core, a multifaceted interaction hub essential for HIV-1 replication.

## Results

We have previously reported numerous HIV-1 inhibitors derived from PF74 and have identified several chemical modifications that improve the antiviral potency of these compounds (29, 60, 63). PF74 contains three aromatic moieties, the “R1” phenyl ring, the “R2” benzyl ring that mimics the phenylalanine of the FG-dipeptide, and the “R3” indole ring (Supplementary Figure S1) (16, 26, 27, 63). By selecting several potent analogs, we have a small library that can be used to probe specific moieties and chemical interactions within the FGBP (60). Selected compounds include the lead compound ZW-1261 as well as ZW-1260 and ZW-1559 that remove the C2-methyl of PF74 R3 and add a C5-hydroxyl, and ZW-1514 and ZW-1517 that modify R3 to have an N-ethyl group (Supplementary Figure S1) (29, 60, 63, 76). Further, ZW-1260 has a *p*-methyl group in R1, while ZW-1261 and ZW-1517 have a *p*-Cl group. While LEN contains the scaffold of the FG-motif (27, 58), it is more complex compared to PF74, thus the PF74-derriviative compounds enable specific chemical probing of the FGBP (60).

It has been established that PF74 has a bimodal mechanism of action depending on the concentration, stabilizing lattice and perturbing core stability (26, 56, 63, 77–79). To assess if a compound can increase or prevent capsid lattice assembly, an *in vitro* assay has been routinely employed to determine the effect of a ligand or condition on CA•CA interactions. Since *in vitro* CA assemblies like tubes and CLPs make a solution turbid, absorbance at 350 nm (A_350_) can be used as a proxy for CA assembly, and thus we are able to infer how a compound or condition can impact the rate of lattice formation (Figure 2A) (10, 72, 80–82). For example, excess NaCl has been used to initiate capsid assembly to determine if compounds increase, decrease, or have no effect on assembly rate; other factors that initiate CA assembly and increased A_350_ are pH <7.0 and the cofactor IP6 (10, 72, 80–82).. For example, the capsid assembly inhibitor peptide (CAI) binds to the CA_CTD_ and prevents CA_HEX_•CA_HEX_ interactions decreasing assembly rates, while the scrambled form of CAI (scCAI) has no effect on assembly (83), and the antiviral PF74 forms interactions between adjacent CA monomers within CA_HEX_ thus increasing assembly rates at pH 8.0 (Figure 2B) (16, 76, 80). We find that like PF74, the other FGBP-targeting antivirals like ZW-1261, ZW-1517 and LEN also increase the rate of CA assembly when induced with excess NaCl (Figure 2C).

**Figure 2:**
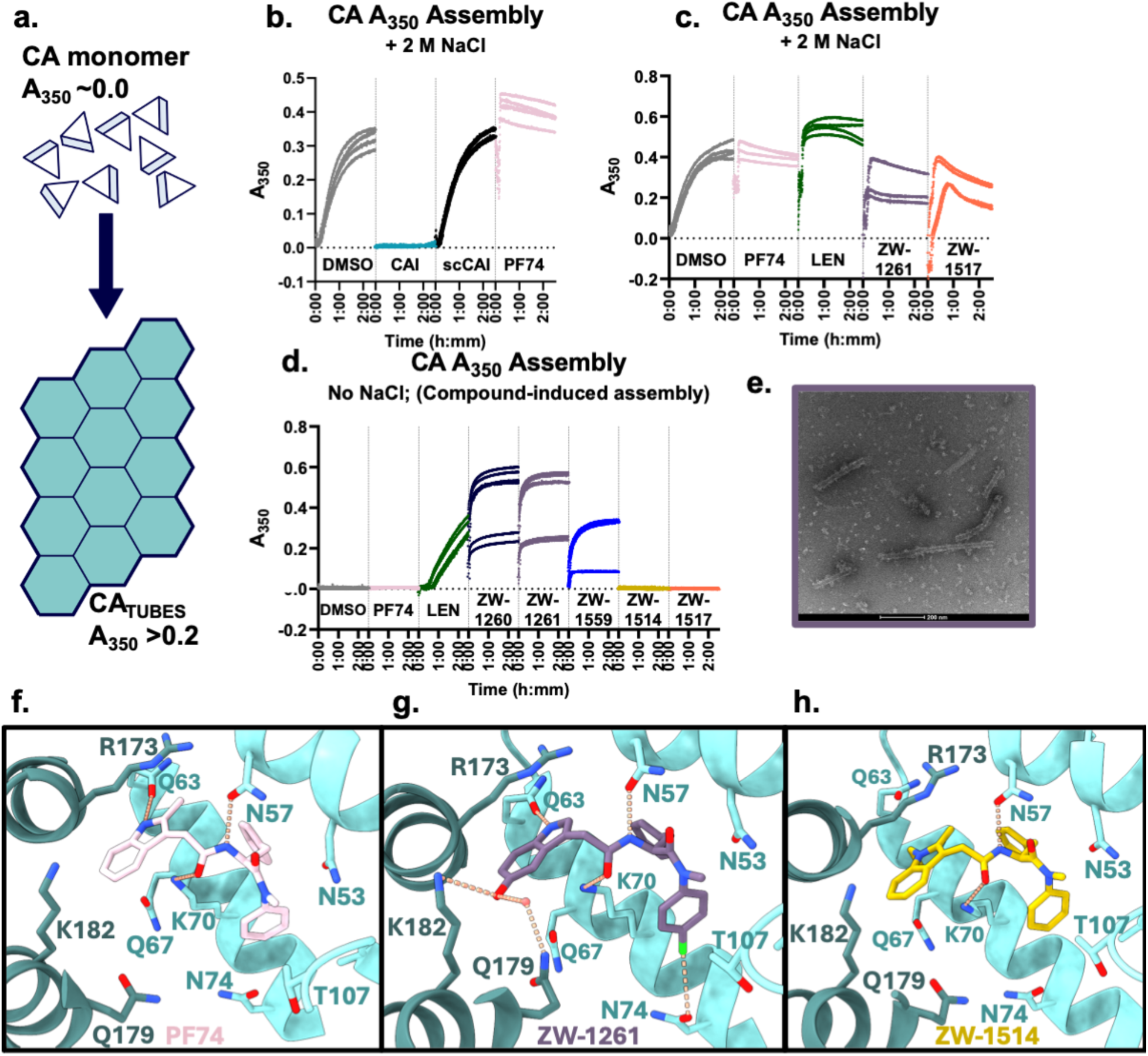
ZW-1261 robustly affects the *in vitro* assembly characteristics of HIV-1 capsid protein (CA), rapidly inducing tubular lattice at pH 8.0. **A.** Overview of *in vitro* assembly assay (76, 81, 82, 84), where soluble CA monomers polymerize into macromolecular assemblies, (Increase in absorbance at 350 nm, A_350_) causing the solution to become opaque. Assembly is initiated with up to 2 M NaCl (**B** & **C**) or with only compounds at 0 M NaCl (**D** & **E**). **B.** Effects of different compounds on the rate of CA assembly, showing inhibition assembly with the CAI peptide (cyan) (83), promotion assembly with PF74 (pink) (76), and having no effect with scrambled CAI peptide (scCAI, black) (83). **C.** The pretreatment of CA with FG-binding compounds increases the rate of *in vitro* assembly when initiated by 2 M NaCl compared to DMSO. **D.** Compounds with the OH-containing indole ring (ZW-1260, ZW-1261, and ZW-1559, see Supplementary Figure S1) spontaneously initiate assembly in the absence of NaCl, even more rapidly than LEN, at CA 50 µM CA and 50 µM compound. **E.** Representative negative-stain TEM image of CA tubes formed by ZW-1261 treatment. Tubes were formed as in D. **F-H.** Comparing the structures of various FG-binding compounds find the R3 indole modifications in ZW-1261 bridge two adjacent monomers within a hexamer with a coordinated water. **F**. PF74 [PDB: 4XFZ (42)]; **G.** ZW-1261 [PDB: 7M9F (13)]; **H.** ZW-1514 [PDB: 9DTM (60, 76)].

Interestingly, we find that some, but not all FGBP-targeting antivirals can initiate lattice assembly in the absence of other assembly-inducers. At 0 M NaCl and pH 8.0 that disfavors CA assembly, adding equimolar ZW-1261 and other compounds with the same R3 indole modifications rapidly create a turbid solution as well as LEN initiating turbidity after an hour of incubation (Figure 2D). We have confirmed by negative stain electron microscopy (EM) that the assembly initiated by ZW-1261 is tubular, indicating a CA_HEX_-only lattice (Figure 2E). The independent induction of assembly by ZW-1261 is distinctly different than the phenotype observed for PF74 and other analogs ZW-1514 and −1517. We hypothesize this is due to the multiple additional contacts that the R3 of ZW-1261 forms with the CA_CTD_, bridging the two monomers that comprise the FGBP pocket with a coordinated water (Figure 2F–2H) (60). The other molecules that induce assembly, ZW-1260 and ZW-1559, have an identical R3 to ZW-1261 (Figure 2D, Supplementary Figure S1). Further, LEN is a large molecule that has been shown to have coordinated waters with the CA_NTD_ but interact with multiple monomers of a CA_HEX_, and thus, it likely initiates assembly in a similar way (10, 58, 73). We find from these experiments that ZW-1261 and other compounds with an R3 that can bridge two adjacent monomers of a CA_HEX_ are powerful assembly-initiating agents.

We have previously used TSAs to identify and quantify the effects of PF74 analogs, and other compounds that target CA on the thermal stability (29, 59, 63, 71, 82). All compounds used in this study increase the 50% melting temperature (ΔT_m_) of soluble CA_HEX_ at pH 8.0 (60, 71, 84); this data differs from previous reports that used 50 mM sodium phosphate buffer (pH 8.0), where this uses 50 mM tris (pH 8.0) to not compete at site 5 (71) (Supplementary Figure S2A). FGBP-compounds differ with LEN being the most thermally-stabilizing with a ΔT_m_ of 18.1 ± 2.0°C, and ZW-1261 and PF74 alone have a ΔT_m_ of 11.4 ± 1.8°C and 8.5 ± 2.5°C respectively (Supplementary Figure S2A). Compounds that bind to the central pore of CA_HEX_ (Site 5), IP6 and hexacarboxybenzene (HCB), are also extremely stabilizing with a ΔT_m_ of 12.4 ± 1.4°C and 13.3 ± 1.3°C (Supplementary Figure S2A). While these compounds alone are stabilizing, we find there is additional stabilization of the CA_HEX_ when ligands that bind to the FGBP and central pore are both present, with IP6+ZW-1261 ΔT_m_ = 20.7 ± 1.3°C, HCB+ZW-1261 ΔT_m_ = 20.3 ± 1.4°C, IP6+LEN ΔT_m_ = 27.9 ± 1.5°C, HCB+LEN ΔT_m_ = 28.8 ± 1.3°C (Supplementary Figure S2A). However, there is no additional stabilization observed if there are multiple compounds that bind to the same pocket, as it appears that the ΔT_m_ is equal to the ΔT_m_ of the most-stabilizing molecule: LEN+ZW-1261 ΔT_m_ = 18.3 ± 1.4°C *vs.* LEN alone ΔT_m_ = 18.1 ± 2.0°C, PF74+ZW-1261 ΔT_m_ = 11.1 ± 1.7°C *vs.* ZW-1261 alone ΔT_m_ = 11.4 ± 1.8°C, or HCB+IP6 ΔT_m_ = 13.0 ± 1.3°C *vs.* HCB alone ΔT_m_ = 13.3 ± 1.3°C (Supplementary Figure S2A).

We aimed to solve the structure of ZW-1261 and IP6 in the context of an assembled CA lattice containing both CA_HEX_ and CA_PENT_. Applying an excess of IP6 to monomeric CA also forms CLPs *in vitro* that are closed containers with 12 CA_PENT_ (Figures 1F and 3A) (49, 66). As mentioned above, treating CA monomers with ZW-1261 alone promote tubes, suggesting CA_HEX_-only assemblies (Figures 1E, 2E, 3B). When IP6 and ZW-1261 are added together simultaneously to monomeric CA, a mixture of elongated but closed tubes and irregularly-shaped assemblies are formed (Figure 3C). If CLPs are first formed with excess IP6 then treated with equimolar ZW-1261, the resulting assemblies appear like broken CLPs with irregular shape (Figure 3D). These were visualized by negative stain TEM and morphologies were quantified by appearance similar to previous reports on CLP morphology (41, 72); we find more elongated and tubular assemblies in the presence of ZW-1261 (Figure 3E).

**Figure 3:**
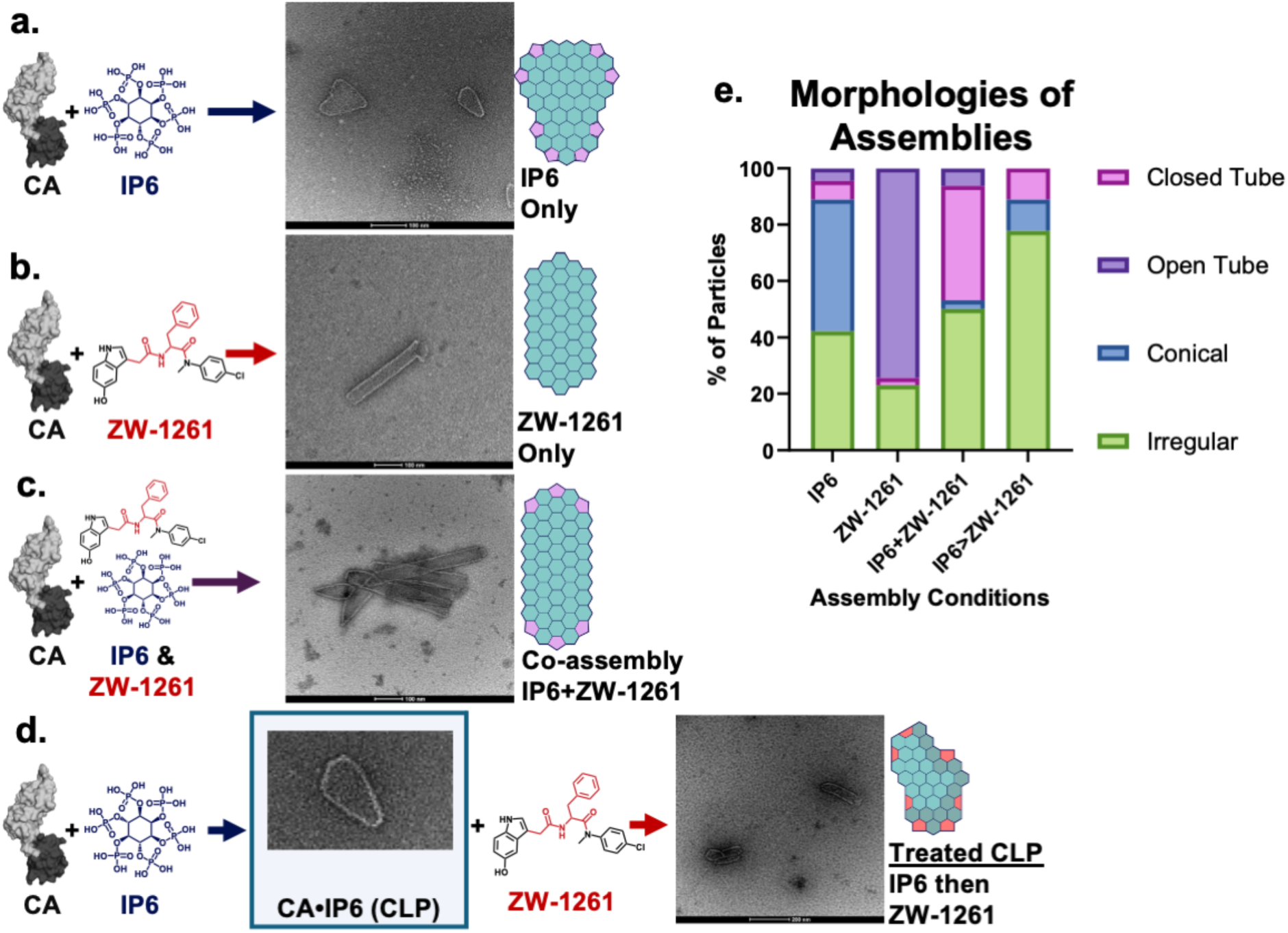
The order-of-addition for compounds that bind CA at different sites impact the resulting macromolecular structures at pH 6.1. **A-D.** Negative-stain transmission EM images of assemblies at pH 6.1. **A.** When excess IP6 is added to purified HIV-1 CA at pH 6.1, CLPs form *in vitro* that resemble native-like cone shapes (see Figure 1F) (49, 66). **B.** When ZW-1261 is added alone, CA form open tubes and aggregates (see Figure 1E). **C.** When ZW-1261 and IP6 are added together simultaneously, elongated cones and oddly-shaped cores form that do not recapitulate the dimensions of an infectious cores. This assembly reaction mimics the maturation phase of the HIV-1 replication cycle in a treated individual, lacking the spatial constraints of the envelope. **D.** When IP6-induced CLPs are then treated with ZW-1261, the lattice aggregate and appear damaged. This assembly reaction mimics treatment of an infectious mature particle. **E.** Quantification of particle morphologies.

To investigate the effects of lattice damage caused by ZW-1261 on the CLPs, we utilized single-particle cryo-EM to solve structures of CA_HEX_ and CA_PENT_ as previously reported for IP6-only assemblies (49, 66). Using this system, after treating the pre-assembled CLPs with equimolar ZW-1261, we were able to resolve a map of CA_HEX_ bound with ZW-1261 along with six-neighboring CA_HEX_. However, unlike other CLP systems, we were unable to resolve a structure of CA_PENT_ despite attempts at manual particle picking at curved interfaces of the damaged CLPs (Figure 4A-D).

**Figure 4:**
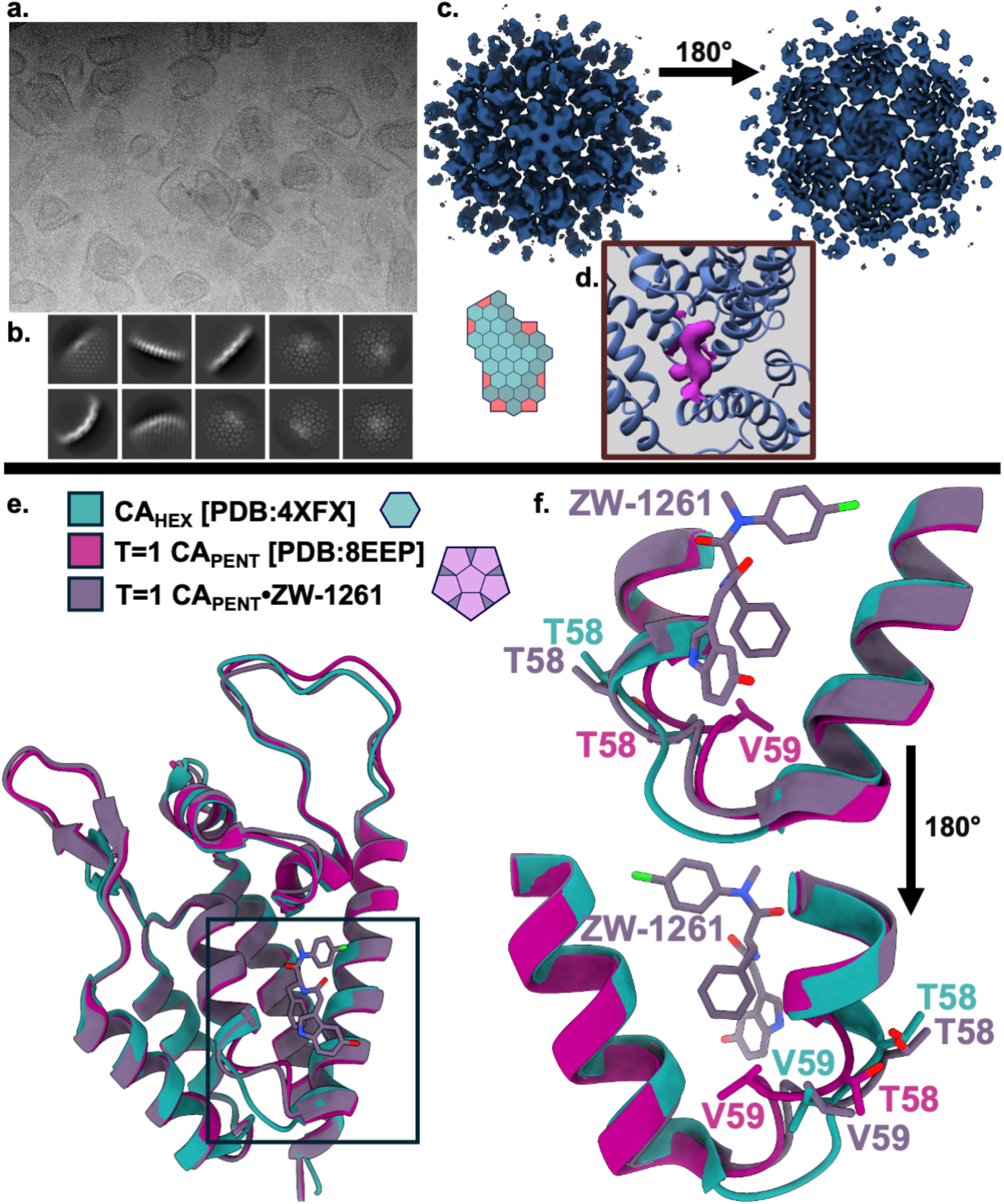
Structural comparisons of assemblies treated with ZW-1261 show morphological and structural differences at CA_PENT_ sites. **A.** Representative micrograph showing soluble CLPs treated with ZW-1261 have a damaged appearance. **B.** The 2D class averages for CLP single-particle cryoEM could not identify classes with CA_PENT_. **C and D.** We were able to solve a map of CA_HEX_ in CLPs at 4.76 Å, though no CA_PENT_ were observed. There is density for ZW-1261 in the FGBP of CA_HEX_ that does not correspond to the docked structure of CA [PDB: 4XFZ (42)]. **E.** We solved these pre-formed T = 1 CA particles treated with ZW-1261 and find the TVGG loop has a mixed conformation (purple). **F.** Cryo-EM structure of ZW-1261 treated T = 1 icosahedrons made of CA[G60A/G61P] (purple) at 2.32 Å resemble both a hexamer (cyan) [PDB: 4XFX (42)] at Thr58 and Val59, and a pentamer (pink) [PDB: 8EEP (41)] at the mutated Ala60 and Pro61 that impose pentamerization (41).

Thus, to resolve the structural interface of CA_PENT_ bound with ZW-1261, we utilized the previously reported T = 1 icosahedral CA_PENT_-only assemblies that have a mutation at G60A and G61P to force the 58-TVGG-61 into a the elongated α3 found in CA_PENT_ that was shown to have a sterically-blocked FGBP (41). The monomeric CA[G60A/G61P] is assembled into icosahedrons with IP6 + 150 mM NaCl (Supplementary Figure 3A). Upon treatment of these icosahedrons with either ZW-1261 or LEN, a minor population of the 20 nm T = 1 icosahedrons appeared larger (Supplementary Figure S3B–S3C). We were able to solve for the T = 1 apo structure at 2.74 Å in a conformation identical to a previously reported (Supplementary Figure S3D) (41).

Once these T = 1 assemblies were treated with ZW-1261, we were able to solve a structure of the CA_PENT_-only icosahedron with compound partially bound to the FGBP at 2.32 Å (Figure 4E). While the mutated residues at G60A/G61P in the loop are forced to be CA_PENT_-like, ZW-1261 treatment causes the preceding Thr58 to a CA_HEX_-like position (Figure 4F). Thr58 is one of the few residues that forms differing interactions in CA_HEX_ *vs.* CA_PENT_ (41, 43), and influences the rotational pitch of α3 that switches the FGBP to be open (in CA_HEX_) or closed (in CA_PENT_); for ZW-1261 to interact at this site, the Met66 cannot be in the CA_PENT_ position that blocks the FGBP. The CA_PENT_-only construct have increased distance between the two monomers that make an FGBP, so unlike the structures of CA_HEX_, both from the CLPs and X-ray crystal structures (13, 60, 76), we only find interactions of ZW-1261 with the CA_NTD_ and not the adjacent CA_CTD_ in the T = 1 icosahedrons (Figure 4F). This is reminiscent of the CA•PF74 structures of CA_NTD_-only crystals (16). Boths solved treated and untreated icosahedrons had two IP6 molecules bound to the central pore. Overall, the mutated CA[G60A/G61P] that forms CA_PENT_ can interact with ZW-1261, despite a theoretically blocked pocket (41), and the compound converts the structure of the five CA monomers of the CA_PENT_ into a CA_HEX_-like conformation.

## Discussion & Conclusions

Previous reports have shown that ZW-1261 is a potent antiretroviral compound that interacts at the FGBP of CA_HEX_ (13, 29, 60, 76). Here, we report the structural impact of ZW-1261 on capsid lattice in the presence of IP6; this assembly cofactor leads to formation of CA_PENT_ and capsid closure (31, 44, 48, 49, 66, 67, 72). We observe that like IP6, ZW-1261 interactions with CA monomers facilitate lattice assembly. However the morphology and apparent integrity of the particles assembled *in vitro* varies with the order of small-molecule addition (Figure 3). We find that ZW-1261 strongly favors CA_HEX_ formation and rapidly induces tubular assemblies, implying a composition of only CA_HEX_. The simultaneous addition of IP6 and ZW-1261, conditions that mimic the late stages of virus production, leads to formation of irregular morphologies and tubes with closed ends, suggesting a combination of both CA_HEX_ and CA_PENT_. This is not the case for stepwise addition, which mimics the early stages of infection; for pre-formed CLPs assembled *in vitro* with IP6, an unconstrained system, ZW-1261 addition leads to damage in the particles and an apparent absence of CA_PENT_. Our findings for ZW-1261 are consistent with reports of LEN and capsid defects (73–75), including one that showed an increase in lattice fractures and a loss of curvature (“declinations”) over time in purified HIV-1 cores following LEN addition (74). This also aligns with the difference in A_350_ rates between ZW-1261 and LEN (Figure 2B).

Only when CA_PENT_ are constrained by reported mutations G60A and G61P (41, 43), we were able to observe ZW-1261 binding to the FGBP of CA_PENT_ (Figure 4). For this occupancy to occur, we find the local conformation of the TVGG loop to mimic the unconstrained CA_HEX_ (41, 43). By using mutations to force CA_PENT_ formation, we can observe what appears to be an intermediate in the structural interconversion between CA_PENT_ and CA_HEX_ that has CA_PENT_ stoichiometry but CA_HEX_-like structure at Thr58 and Val59.

Further, it was previously shown that Met66 pivots into the FGBP when the TVGG is in the CA_PENT_-like extended loop conformation, which would sterically block the binding of FG-containing host factors like Nup153 (41, 43, 85). The M66I mutation imposes slight resistance to ZW-1261 and strong resistance to LEN, and since Met66 was shown to change positions with the TVGG switch (41, 43, 64, 85, 86), it is possible that the mutation affects the equilibrium between CA_PENT_ to CA_HEX_. Notably, the M66A mutation leads to formation of CA_PENT_-only T = 1 icosahedrons that are converted to tubes upon binding of LEN to the FGBP (41, 73). Combining occupancy at the FGBP with the IP6 causes a strong increase in CA_HEX_ thermal stability, aligning with previous reports of increased lattice stability but does not necessarily improve the integrity of the core (56, 60, 73, 78).

Of note, the distance between monomers of the CA_PENT_ that build the T = 1 icosahedron are increased compared to a CA_HEX_, and as such, we only observe interactions of ZW-1261 with the CA_NTD_. While the FGBP pocket is typically formed between the CA_NTD_ and an adjacent CA_CTD_, inhibitor binding to a construct of only a CA_NTD_ has been previously reported for PF74, however, the R3 group of PF74 differs between the CA_NTD_ and CA_HEX_ structures (26, 27, 42); we observe only partial density of ZW-1261 R3, consistent with multiple orientations of the indole ring that due to the loss of interactions with the CA_CTD_ as previously seen in crystallographic structures (13, 60, 76).

Overall, ZW-1261 is a potent inducer of CA_HEX_ lattice formation by bridging two monomers through its R3 hydroxyl group (60, 76) (Figure 2G), and its binding to CA_PENT_ requires structural rearrangements likely breaking an unconstrained core lattice (Figure 4). This is in opposition to the reported effects of IP6 and other polyanions that induce curvature that forms CA_PENT_ (31, 44, 49, 66). We see that both ZW-1261 and IP6 can bind CA_HEX_ simultaneously, as seen in resolved structures that were verified by TSAs and nMS (Supplementary Figure S2). ZW-1261 binding to the FGBP interferes with conical core formation, even in the presence of IP6 (Figures 3C & 4A), and changes the morphology of the resulting assemblies. Collectively, this suggests a mechanism for inhibitor binding the FGBP that impacts capsid closure and core integrity at the CA_PENT_.

## Materials

Lenacapavir (LEN) was purchased from MedChemExpress (Monmouth Junction, NJ). Synthesis of PF74, ZW-1260, ZW-1261, and ZW-1559 were reported previously (29). Synthesis of ZW-1514 and ZW-1517 were reported previously (63). All antiviral compounds were suspended in ≥99.9% DMSO (Sigma-Aldrich; St. Louis, MO). Phytic Acid (IP6) 50% in H_2_O was purchased from TCI America (Portland, OR). Mellitic Acid (Hexacarboxybenzene, HCB) was purchased from Sigma-Aldrich (St. Louis, MO) and suspended in dH_2_O. CAI and scCAI sequences (83) were synthesized by Genscript (Piscataway, NJ).

PDB models used include WT CA monomers [PDB: 4XFX (42)] and crystallographic p6 symmetry of 4XFX is used for representing CA hexamers, CA pentamers [PDB: 3P05 (87)], fullerene-cone mature capsid assemblies [PDB: 3J3Q (88)], CA Tubes [PDB: 3J4F (88)], and T = 1 icosahedrons [PDB: 8EEP (41)]. For the T = 4 icosahedron, 4XFX (42) hexamers and 3P05 (87) pentamers were docked into the T = 4 map [EMD-9733(50)] *via* ChimeraX (51, 89). Models of WT CA treated with PF74 [PDB: 4XFZ (42)], ZW-1261 [PDB: 7M9F (13)], and ZW-1514 [PDB: 9DTM (60, 76)] were shown in Figure 1. The crystal structures of WT CA and ZW-1261 [PDB: 7M9F (13)] or T = 1 icosahedrons [PDB: 8EEP (41)] were used for model building. Structures were visualized in ChimeraX (89).

## Methods

### Expression and Purification of HIV-1 Capsid (CA)

WT monomeric HIV-1 capsid protein (CA) was cloned in a pET11a expression plasmid, provided by Dr. Chun Tang (Peking University). For crystallography experiments, CA was overexpressed in BL21(DE3)RIL *E. coli* and CA was purified by ammonium sulfate precipitation followed by anion exchange chromatography and stored in 20 mM Tris (pH 8.2) with 40 mM NaCl as previously described (42). For mature capsid assemblies and electron microscopy, monomeric CA was overexpressed in NiCo21(DE3) *E. coli* and purified by ammonium sulfate precipitation followed by desalting chromatography (HiPrep), subtractive ionic chromatography, and size-exclusion chromatography (SEC, Superdex 200 10/30 GL), flash-frozen and stored in 25 mM Tris (pH 8.0) as previously described (49).

Cross-linked CA hexamers, containing A14C/E45C/W184A/M185A mutations for disulfide stabilization (CA121 or CA_HEX_), were cloned into a baterial expression plasmid, provided by Dr. Owen Pornillos (University of Utah) (38). *E. coli* BL21(DE3)RIL was used for overexpression and CA_HEX_ was purified as previously described (38, 59), with additional size-exclusion chromatography step for added protein purity to remove non-crosslinked CA (HiLoad™ 26/600 Superdex 200) in storage buffer (20 mM Tris pH 8.2 and 40 mM NaCl).

Constitutive pentameric CA, containing G60A/G61P mutations for obligate pentamerization (41), was cloned into a pET11a vector (Genscript; Piscataway, NJ) with C-terminal 6xHIS-Tag. This protocol was adapted from (90). Overexpression was induced with 100 µM IPTG in NiCo21(DE3) *E. coli* at 18°C overnight. Bacteria were lysed by sonication (8 min total, 30 s on, 30 s off, 75% amplitude) in 50 mM Tris (pH 8.0) with 1 mM TCEP and 300 mM NaCl. The soluble lysate fraction was incubated with nickel (Ni^2+^) beads for >1 h at 4°C, washed with lysis buffer, and eluted with 20-100 mM imidazole. Unassembled CA[G60A/G61P] was concentrated to ∼25 mg/mL and dialyzed into 25 mM MES (pH 6.0) with 1 mM TCEP and 50 mM NaCl before freezing.

### A_350_ *in vitro* CA Assembly Assay

The CA assembly assay was modified from a previously described method (49, 81, 82, 84). 100 µM of CA monomers (2X solution) was prepared in 50 mM Tris (pH 8.0). For testing NaCl-induced assembly, the 2X CA solution was treated with equimolar compound (100 μM, ≤2% DMSO) on ice for ∼30 min. These 2X Solutions were dispensed into a 96-well plate and mixed 1:1 with 4 M NaCl in 50 mM Tris (pH 8.0) to initiate assembly. For testing compound-only assembly, 2X solutions of antivirals were made separately and mixed 1:1 in the plate to initiate assembly.

Absorbance at 350 nm (A_350_) was measured every 25 seconds for 150 minutes at room temperature with a Synergy Neo 2 (BioTek) plate reader. Samples containing the 1X solution of CA, compound, and 2 M NaCl were background subtracted from a blank well that lacked NaCl. In the compound-only assembly assay, controls that lacked CA were used for background subtraction. GraphPad Prism 10 was used for visualization and statistical analysis.

### Thermal Shift Assay (TSA)

7.5 µM CA121_HEX_ was incubated with 20 µM of each compound (≤1% DMSO) in 50 mM Tris Buffer (pH 8.0) (71). Following a 30 minute incubation on ice, samples were then mixed with dye to a final concentration of 1X SYPRO™ Orange dye in a qPCR plate and samples were heated from 25–95°C QuantStudio 3 Real-Time PCR Systems (Thermo Fisher Scientific) as previously described (29, 82, 84). Thermal profiles were analyzed with Protein Thermal Shift Software v1.3 (Applied Biosystems) and visualized with TSAR (71). Statistical significance was determined by comparing the treated condition to the DMSO vehicle with a two-sided unpaired t-test in R.

### Capsid Assemblies & Negative Stain Transmission Electron Microscopy (TEM)

Capsid-like particles (CLPs) and other WT assemblies were formed by mixing WT CA monomers (250 μM CA in 50 mM MES (pH 6.2)) with equimolar ZW-1261, 5X excess IP6:CA, or a mix of both 37°C. Protocol adapted from (49, 66). Samples were diluted 1:20 before spotting on the grid and stained with 0.75 % Uranyl Formate or Acetate, then imaged on an FEI Talos 120 KV with LaB6 and 4k Ceta detector (ThermoFisher). Assembly appearance was quantified similar to previous reports on CLP morphology (41, 72).

For T = 1 icosahedrons, assembly occurred in a final condition of 50 mM MES (pH 6.0), 50 mM NaCl, 5 mM IP6 and 1 mM TCEP at 37°C for 2 h prior to purification by SEC (Superdex 200 10/30 GL) in 25 mM Tris (8.0), 150 mM NaCl, 0.5 mM IP6 and 0.5 mM TCEP. Protocol adapted from (34, 41). Samples were diluted 1:5 before spotting on the grid and stained with 0.75 % Uranyl Formate or Acetate, then imaged on an FEI Talos 120 KV with LaB6 and 4k Ceta detector (ThermoFisher).

### Cryo-EM grid preparation

Purified T = 1 icosahedral spheres (T1s) were assembled as described above (41, 90), and pooled to a concentration of ∼10 µM. For treated icosahedrons, the assemblies were incubated with a molar excess of ZW-1261 at 4°C for 1 hour prior to blotting. UltrAufoil R 2/2, 200 mesh, Au grids were glow discharged and 3 µL of sample was applied to the grids followed by blotting and vitrification using a Vitrobot at 4°C and 80% humidity. Untreated T1s followed a similar protocol, however, UltrAufoil 1.2/1.3, 300 mesh, Au grids were used.

CLPs were prepared as described above (34, 41), and ZW-1261 was added at a 1:1 molar ratio of CA:compound for 10 minutes at 37°C. Samples were diluted 1:4 before applying the sample to the grid. Quantfoil R 2/2, 200 mesh, Cu grids were glow discharged and 3 µL of sample was applied to the grids followed by blotting and vitrification using a Vitrobot.

### Single Particle Cryo-EM data processing

For both treated and untreated T1s, initial processing of motion correction was done in Relion 4.0 (91). Motion corrected micrographs were then transferred to CryoSPARC for further processing (92). Patch CTF estimation was performed followed and micrographs were curated. Particles were picked using blob picker with diameter of 50 – 300 Å. Using a box size of 480 pixels, selected particles were extracted and 2D classifications were determined. After iterative refinement of the particle stack using 2D classification, a series of homogeneous, heterogenous and non-uniform refinements were performed. Global and local CTF corrections were performed followed by a final round of non-uniform refinements. For the treated icosahedrons, Relion was used for 3D refinement and Bayesian polishing (91). A similar pipeline was performed for CLPs for initial processing in Relion and CryoSPARC, however the lack of CA_PENT_ led to manual particle picking at curved interfaces of CLPs. See Supplementary Table S1 for collection and processing details.

### Native Mass Spectrometry (nMS)

CA121 (38) samples were buffer exchanged into 200 mM ammonium acetate (Sigma Aldrich) using micro P6 spin columns (Bio-Rad), and stored at 4°C overnight before analysis. 10 mM IP6 in water was prepared fresh each day from 1.1 M stock, adjusting the pH to 6. All other ligands were prepared in DMSO. For all nMS experiments CA121 was diluted to approximately 6 µM monomer concentration, however, a low intensity species at ∼18 kDa is observed in the samples and therefore exact CA121 concentration cannot be reported. For ligand binding experiments CA121 was diluted to 6 µM monomer concentration, ligands were added at 1, 3, 6, 9, and 18 µM concentration. DMSO, which is known to reduce the average charge states in nMS at low concentrations (93), was added to the apo protein and IP6 samples at 4% by volume to match conditions used for ligands resuspended in DMSO. Samples were incubated on ice for a minimum of 15 minutes prior to analysis.

All experiments were performed on a Q Exactive ultra high mass range (UHMR) Orbitrap (Thermo Fisher Scientific). Samples were introduced into the MS using nano-electrospray ionization, pulled in-house using a P-97 micropipette puller (Sutter instruments). The MS was operated in positive mode, with a capillary temperature of 250 °C, a resolution setting of 6, 250, in-source trapping −30 to −50 V, low detector mode, trap gas 5 (94, 95).

## Acknowledgments

This research was supported in part by the National Institutes of Health (U54 AI170855 to K.A.K., S.R.H., R.A.D., and S.G.S.; R01 AI120860 to Z.W. and S.G.S.; and P30 AI050409 to S.G.S.; R21 AI189247 to K.A.K.; RM1 GM149374 to V.H.W.; F31 AI174951 to W.M.M.) W.M.M. and Z.C.L. were supported in part by T32 GM135060. Electron Microscopy was carried out by Emory University Robert P. Apkarian Integrated Electron Microscopy Core Facility (RRID: SCR_023537). The content is solely the responsibility of the authors and does not necessarily represent the official views of the National Institutes of Health. Additionally, S.G.S. acknowledges funding from the Nahmias-Schinazi Distinguished Chair in Research.

---

**Supplementary Figure S1:**
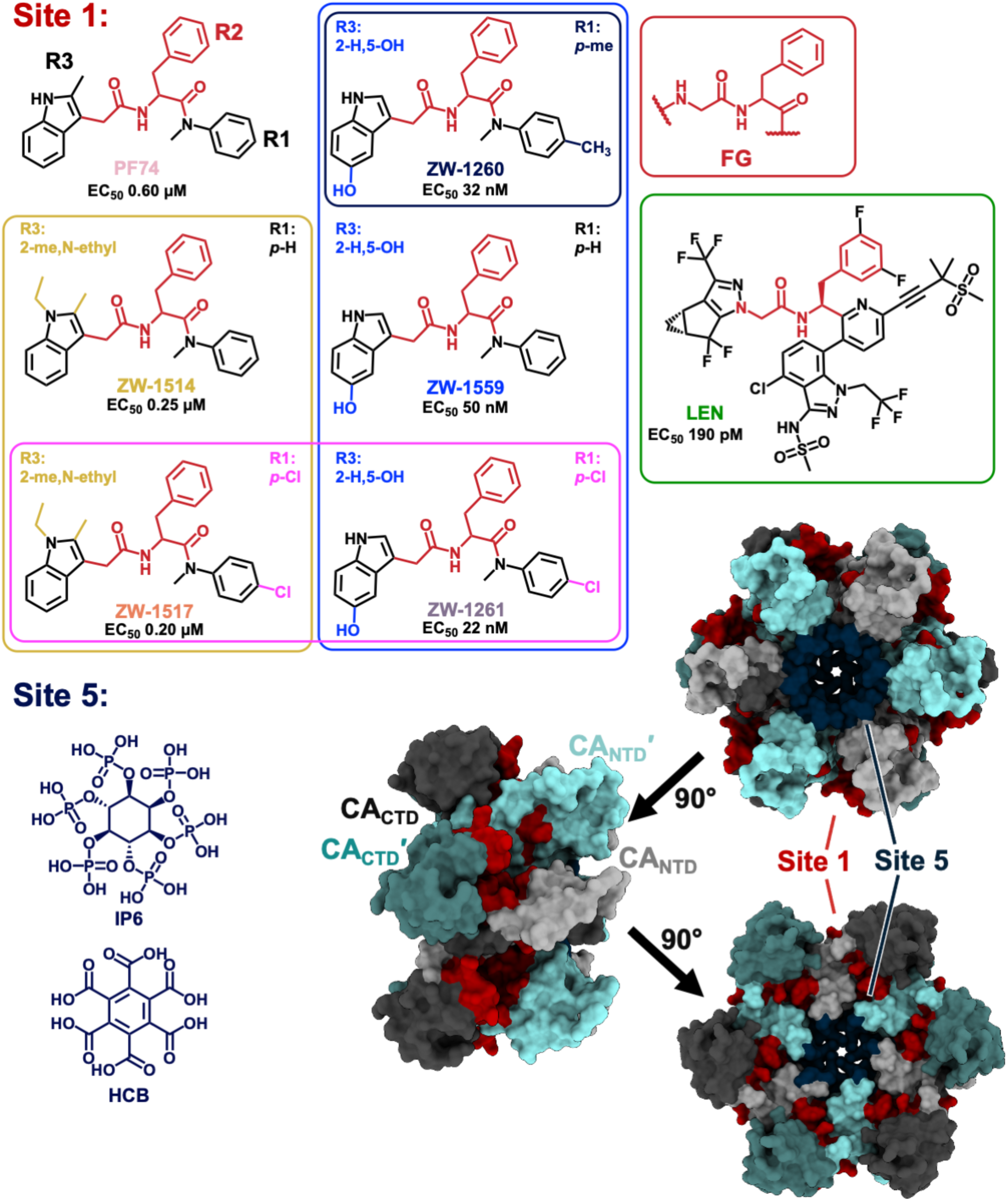
Phenylalanine-Glycine (FG) analog-containing antivirals and other compounds used in this study bind to CA Site 1 or Site 5. PF74 (Pink) was the first compound reported to bind at this site (16) and has three rings (R1, R2, and R3). ZW-1514 (Yellow) and ZW-1517 (Orange) modify the R3 ring with an ethyl group adjacent the methyl. ZW-1559 (Blue), ZW-1260 (Navy Blue), and ZW-1261 (Purple) modify the R3 indole to remove the methyl group on one side and add an alcohol group to the other ring in the indole. ZW-1260, ZW-1261, and ZW-1517 have an additional modification on the R1 ring, with ZW-1260 having a 4-methyl group while ZW-1261 and ZW-1517 have a 4-Cl. Lenacapavir (LEN, Green) is the only approved drug that targets the same site (4). Structures and EC_50_ values from (10, 14, 16, 29, 63). These compounds and others with the FG-scaffold may bind to the FG-binding pocket shared by neighboring protomers in HIV-1 CA hexamers, shaded in red; the central pore for IP6 binding is in navy [PDB: 4XFX (42)]. CA monomers are depicted with the N-terminal domain (NTD) in lighter shades and the C-terminal domain (CTD) in darker shades. IP6 and hexacarboxybenzene (HCB) are shown below that bind site 5, the central pore.

**Supplementary Figure S2:**
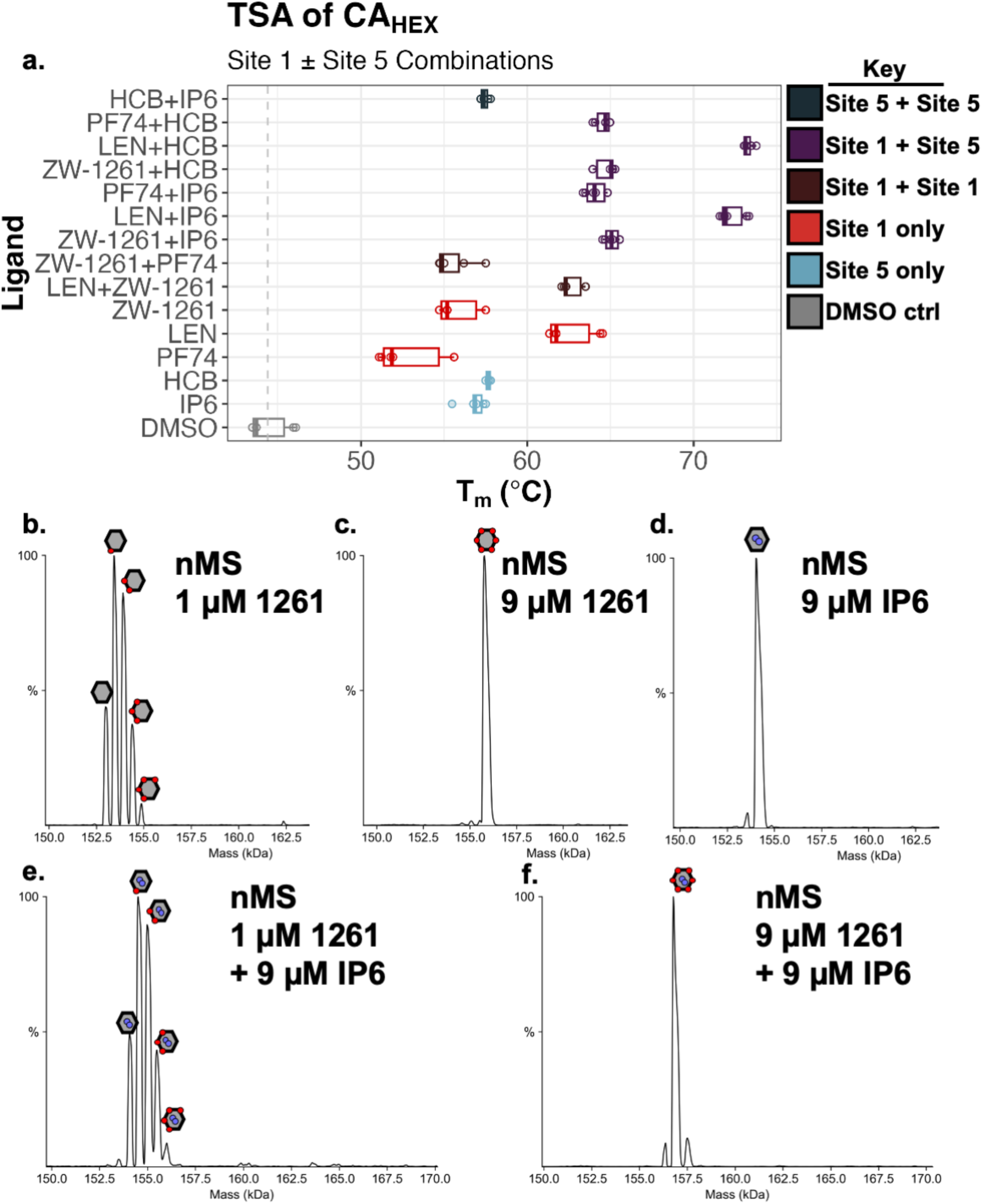
FG-binding pocket can be saturated with one compound, and is further stabilized by IP6. **A.** TSA of CA_HEX_ treated with FG-binding compounds ± Pore-binding compounds. The two different sites have additive effect on the ΔT_m_ for CA_HEX_. Boxplot produced with TSAR (71). **B-F.** Native Mass Spectrometry (nMS) analysis of CA_HEX_ finds ZW-1261 **B.** can bind multiple pockets at 1 µM and **C.** fully saturate a CA_HEX_ by binding to all six pockets at 9 µM. **D.** For a CA_HEX_, two IP6 molecules bind to the central pore at 9 µM IP6. **E**. A combination of 1 µM ZW-1261 and 9 µM IP6 find two IP6 bound and between one and four ZW-1261 bound to a CA_HEX_. **F.** Both IP6 and ZW-1261 can saturate a CA_HEX_ at 9 µM IP6 + 9 µM ZW-1261.

**Supplementary Figure S3:**
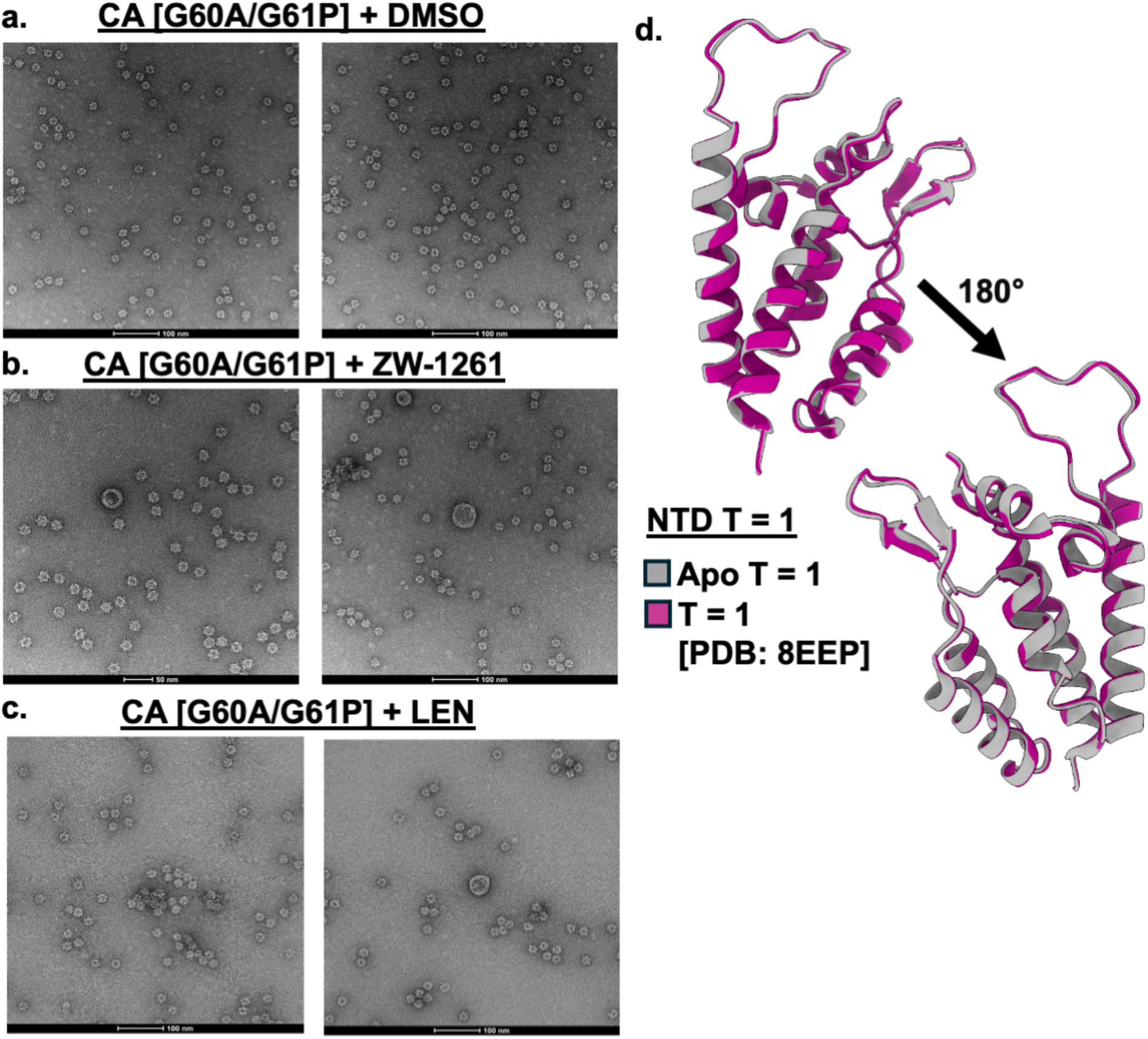
Negative stain TEM images of T = 1 particles treated with FGBP-targeting antivirals. **A.** A pentameric-only assembly of CA forms with mutations G60A/G61P that change the TVGG switch and make a particle with 12 pentamers and 20 nm diameter (41). **B.** Treating pre-formed T = 1 CA particles from (**A**) with ZW-1261 leads to increased aggregation as well as occasionally larger or damaged assemblies. **C.** Same as (**B**), treating T = 1 CA with LEN leads to leads to apparent differences in morphology. **D.** We solved the T = 1 Icosahedral CA_PENT_-only construct in the absence of inhibitor (grey) at 2.74 Å and find the TVGG loop in the untreated condition is identical to the previously reported structure (pink) [PDB: 8EEP (41)]. Thus, adding a HIS-tag to the CA_CTD_ does not impact the FGBP.

**Supplementary Table S1:**
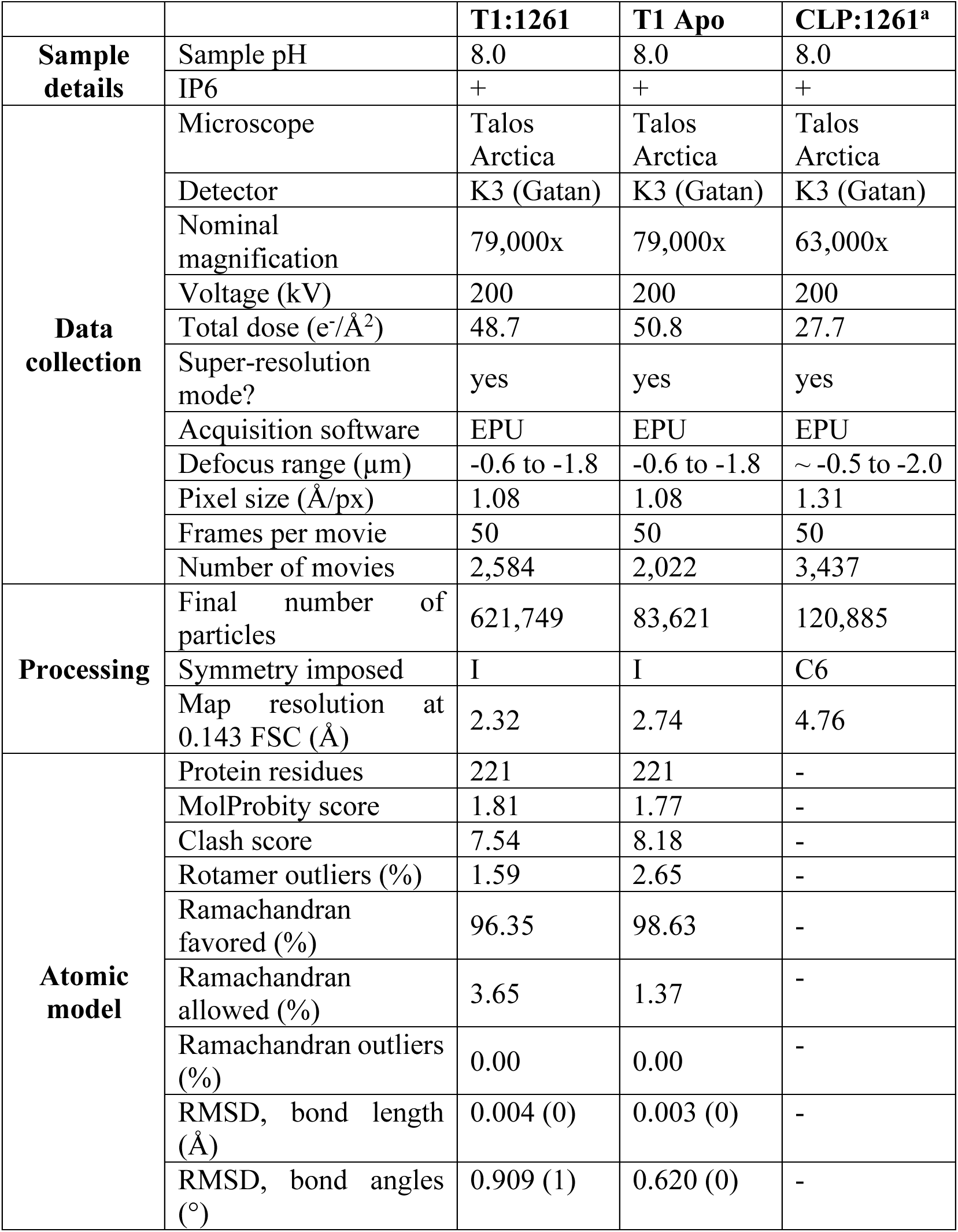
Summary of Cryo-EM Experiments.

## References

1. Ravichandran SM, McFadden WM, Snyder AA, Sarafianos SG. State of the ART (antiretroviral therapy): Long-acting HIV-1 therapeutics. Global Health & Medicine. 2024;6(5):285–94.

2. Sever B, Otsuka M, Fujita M, Ciftci H. A Review of FDA-Approved Anti-HIV-1 Drugs, Anti-Gag Compounds, and Potential Strategies for HIV-1 Eradication. International Journal of Molecular Sciences. 2024;25(7):3659.

3. Thoueille P, Choong E, Cavassini M, Buclin T, Decosterd LA. Long-acting antiretrovirals: a new era for the management and prevention of HIV infection. Journal of Antimicrobial Chemotherapy. 2022;77(2):290–302.

4. McFadden WM, Faerch M, Kirby KA, Dick RA, Torbett BE, Sarafianos SG. Considerations for capsid-targeting antiretrovirals in pre-exposure prophylaxis. Trends in Molecular Medicine. 2025;31(9):801–13.

5. Cilento ME, Kirby KA, Sarafianos SG. Avoiding Drug Resistance in HIV Reverse Transcriptase. Chem Rev. 2021;121(6):3271–96.

6. Grant RM, Liegler T, Defechereux P, Kashuba AD, Taylor D, Abdel-Mohsen M, et al. Drug resistance and plasma viral RNA level after ineffective use of oral pre-exposure prophylaxis in women. AIDS. 2015;29(3):331–7.

7. Himmel DM, Arnold E. Non-Nucleoside Reverse Transcriptase Inhibitors Join Forces with Integrase Inhibitors to Combat HIV. Pharmaceuticals (Basel). 2020;13(6):122.

8. Joint United Nations Programme on HIV/AIDS. The urgency of now: AIDS at a crossroads. Geneva, Switzerland 2024.

9. Bekker LG, Das M, Abdool Karim Q, Ahmed K, Batting J, Brumskine W, et al. Twice-Yearly Lenacapavir or Daily F/TAF for HIV Prevention in Cisgender Women. N Engl J Med. 2024;391(13):1179–92.

10. Link JO, Rhee MS, Tse WC, Zheng J, Somoza JR, Rowe W, et al. Clinical targeting of HIV capsid protein with a long-acting small molecule. Nature. 2020;584(7822):614–8.

11. Ogbuagu O, Segal-Maurer S, Ratanasuwan W, Avihingsanon A, Brinson C, Workowski K, et al. Efficacy and safety of the novel capsid inhibitor lenacapavir to treat multidrug-resistant HIV: week 52 results of a phase 2/3 trial. The Lancet HIV. 2023;10(8):e497–e505.

12. Paik J. Lenacapavir: First Approval. Drugs. 2022;82(14):1499–504.

13. Sun Q, Levy RM, Kirby KA, Wang Z, Sarafianos SG, Deng N. Molecular Dynamics Free Energy Simulations Reveal the Mechanism for the Antiviral Resistance of the M66I HIV-1 Capsid Mutation. Viruses. 2021;13(5):920.

14. Jamey N, Nirosh Udayanga DM, Zhang H, Ravichandran SM, Cai X, Lorson ZC, et al. Design, synthesis and profiling of highly potent antivirals targeting emerging drug-resistant HIV-1 variants. bioRxiv. 2025:2025.12.29.696941.

15. Subramanian R, Tang J, Zheng J, Lu B, Wang K, Yant SR, et al. Lenacapavir: A Novel, Potent, and Selective First-in-Class Inhibitor of HIV-1 Capsid Function Exhibits Optimal Pharmacokinetic Properties for a Long-Acting Injectable Antiretroviral Agent. Molecular Pharmaceutics. 2023;20(12):6213–25.

16. Blair WS, Pickford C, Irving SL, Brown DG, Anderson M, Bazin R, et al. HIV capsid is a tractable target for small molecule therapeutic intervention. PLOS Pathogens. 2010;6(12):e1001220.

17. Rihn SJ, Wilson SJ, Loman NJ, Alim M, Bakker SE, Bhella D, et al. Extreme Genetic Fragility of the HIV-1 Capsid. PLOS Pathogens. 2013;9(6):e1003461.

18. Troyano-Hernáez P, Reinosa R, Holguín Á. HIV Capsid Protein Genetic Diversity Across HIV-1 Variants and Impact on New Capsid-Inhibitor Lenacapavir. Frontiers in Microbiology. 2022;13:854974.

19. Gres AT, Kirby KA, McFadden WM, Du H, Liu D, Xu C, et al. Multidisciplinary studies with mutated HIV-1 capsid proteins reveal structural mechanisms of lattice stabilization. Nature Communications. 2023;14(1):5614.

20. Zila V, Müller TG, Müller B, Kräusslich H-G. HIV-1 capsid is the key orchestrator of early viral replication. PLOS Pathogens. 2022;17(12):e1010109.

21. Hulme Amy E, Kelley Z, Okocha Eneniziaogochukwu A, Hope Thomas J. Identification of Capsid Mutations That Alter the Rate of HIV-1 Uncoating in Infected Cells. Journal of Virology. 2014;89(1):643–51.

22. Burdick RC, Li C, Munshi M, Rawson JMO, Nagashima K, Hu W-S, et al. HIV-1 uncoats in the nucleus near sites of integration. Proceedings of the National Academy of Sciences. 2020;117(10):5486–93.

23. Scott TM, Arnold LM, Powers JA, McCann DA, Rowe AB, Christensen DE, et al. Cell-free assays reveal that the HIV-1 capsid protects reverse transcripts from cGAS immune sensing. PLOS Pathogens. 2025;21(1):e1012206.

24. Jennings J, Shi J, Varadarajan J, Jamieson Parker J, Aiken C. The Host Cell Metabolite Inositol Hexakisphosphate Promotes Efficient Endogenous HIV-1 Reverse Transcription by Stabilizing the Viral Capsid. mBio. 2020;11(6):10–1128.

25. Christensen DE, Ganser-Pornillos BK, Johnson JS, Pornillos O, Sundquist WI. Reconstitution and visualization of HIV-1 capsid-dependent replication and integration in vitro. Science. 2020;370(6513):eabc8420.

26. Bhattacharya A, Alam SL, Fricke T, Zadrozny K, Sedzicki J, Taylor AB, et al. Structural basis of HIV-1 capsid recognition by PF74 and CPSF6. Proceedings of the National Academy of Sciences. 2014;111(52):18625–30.

27. McFadden WM, Snyder AA, Kirby KA, Tedbury PR, Raj M, Wang Z, et al. Rotten to the core: antivirals targeting the HIV-1 capsid core. Retrovirology. 2021;18(1):41.

28. Faysal KMR, Walsh JC, Renner N, Márquez CL, Shah VB, Tuckwell AJ, et al. Pharmacologic hyperstabilisation of the HIV-1 capsid lattice induces capsid failure. eLife. 2024;13:e83605.

29. Wang L, Casey MC, Vernekar SKV, Sahani RL, Kankanala J, Kirby KA, et al. Novel HIV-1 capsid-targeting small molecules of the PF74 binding site. European Journal of Medicinal Chemistry. 2020;204:112626.

30. Kreysing JP, Heidari M, Zila V, Cruz-León S, Obarska-Kosinska A, Laketa V, et al. Passage of the HIV capsid cracks the nuclear pore. Cell. 2025;188(4):930–43.e21.

31. Hudait A, Voth GA. HIV-1 capsid shape, orientation, and entropic elasticity regulate translocation into the nuclear pore complex. Proceedings of the National Academy of Sciences. 2024;121(4):e2313737121.

32. Xu C, Fischer DK, Rankovic S, Li W, Dick RA, Runge B, et al. Permeability of the HIV-1 capsid to metabolites modulates viral DNA synthesis. PLOS Biology. 2020;18(12):e3001015.

33. Stephens C, Naghavi MH. The host cytoskeleton: a key regulator of early HIV-1 infection. The FEBS Journal. 2024;291(9):1835–48.

34. Fu L, Cheng S, Riedel D, Kopecny L, Schuh M, Görlich D. Governed by surface amino acid composition: HIV capsid passage through the NPC barrier. bioRxiv. 2025:2025.03.13.643050.

35. Hou Z, Shen Y, Fronik S, Shen J, Shi J, Xu J, et al. HIV-1 nuclear import is selective and depends on both capsid elasticity and nuclear pore adaptability. Nature Microbiology. 2025;10(8):1868–85.

36. von Schwedler Uta K, Stray Kirsten M, Garrus Jennifer E, Sundquist Wesley I. Functional Surfaces of the Human Immunodeficiency Virus Type 1 Capsid Protein. Journal of Virology. 2003;77(9):5439–50.

37. Byeon I-JL, Meng X, Jung J, Zhao G, Yang R, Ahn J, et al. Structural Convergence between Cryo-EM and NMR Reveals Intersubunit Interactions Critical for HIV-1 Capsid Function. Cell. 2009;139(4):780–90.

38. Pornillos O, Ganser-Pornillos BK, Banumathi S, Hua Y, Yeager M. Disulfide bond stabilization of the hexameric capsomer of human immunodeficiency virus. J Mol Biol. 2010;401(5):985–95.

39. Briggs JAG, Simon MN, Gross I, Kräusslich H-G, Fuller SD, Vogt VM, et al. The stoichiometry of Gag protein in HIV-1. Nature Structural & Molecular Biology. 2004;11(7):672–5.

40. Grime JMA, Dama JF, Ganser-Pornillos BK, Woodward CL, Jensen GJ, Yeager M, et al. Coarse-grained simulation reveals key features of HIV-1 capsid self-assembly. Nature Communications. 2016;7(1):11568.

41. Schirra RT, dos Santos NFB, Zadrozny KK, Kucharska I, Ganser-Pornillos BK, Pornillos O. A molecular switch modulates assembly and host factor binding of the HIV-1 capsid. Nature Structural & Molecular Biology. 2023;30(3):383–90.

42. Gres AT, Kirby KA, KewalRamani VN, Tanner JJ, Pornillos O, Sarafianos SG. X-ray crystal structures of native HIV-1 capsid protein reveal conformational variability. Science. 2015;349(6243):99–103.

43. Stacey JCV, Tan A, Lu JM, James LC, Dick RA, Briggs JAG. Two structural switches in HIV-1 capsid regulate capsid curvature and host factor binding. Proceedings of the National Academy of Sciences. 2023;120(16):e2220557120.

44. Gupta M, Hudait A, Yeager M, Voth GA. Kinetic implications of IP6 anion binding on the molecular switch of HIV-1 capsid assembly. Science Advances. 2025;11(16):eadt7818.

45. Li S, Hill CP, Sundquist WI, Finch JT. Image reconstructions of helical assemblies of the HIV-1 CA protein. Nature. 2000;407(6802):409–13.

46. Lu M, Russell RW, Bryer AJ, Quinn CM, Hou G, Zhang H, et al. Atomic-resolution structure of HIV-1 capsid tubes by magic-angle spinning NMR. Nat Struct Mol Biol. 2020;27(9):863–9.

47. Ganser-Pornillos BK, Yeager M, Pornillos O. Assembly and architecture of HIV. Viral molecular machines. 2011;726:441–65.

48. Mattei S, Glass B, Hagen WJ, Krausslich HG, Briggs JA. The structure and flexibility of conical HIV-1 capsids determined within intact virions. Science. 2016;354(6318):1434–7.

49. Highland CM, Tan A, Ricaña CL, Briggs JAG, Dick RA. Structural insights into HIV-1 polyanion-dependent capsid lattice formation revealed by single particle cryo-EM. Proceedings of the National Academy of Sciences. 2023;120(18):e2220545120.

50. Zhang Z, He M, Bai S, Zhang F, Jiang J, Zheng Q, et al. T = 4 Icosahedral HIV-1 Capsid As an Immunogenic Vector for HIV-1 V3 Loop Epitope Display. Viruses. 2018;10(12):667.

51. Lau D, Walsh JC, Mousapasandi A, Ariotti N, Shah VB, Turville S, et al. Self-Assembly of Fluorescent HIV Capsid Spheres for Detection of Capsid Binders. Langmuir. 2020;36(13):3624–32.

52. Matreyek KA, Yücel SS, Li X, Engelman A. Nucleoporin NUP153 Phenylalanine-Glycine Motifs Engage a Common Binding Pocket within the HIV-1 Capsid Protein to Mediate Lentiviral Infectivity. PLOS Pathogens. 2013;9(10):e1003693.

53. Shen Q, Kumari S, Xu C, Jang S, Shi J, Burdick RC, et al. The capsid lattice engages a bipartite NUP153 motif to mediate nuclear entry of HIV-1 cores. Proceedings of the National Academy of Sciences. 2023;120(13):e2202815120.

54. Dickson CF, Hertel S, Tuckwell AJ, Li N, Ruan J, Al-Izzi SC, et al. The HIV capsid mimics karyopherin engagement of FG-nucleoporins. Nature. 2024;626(8000):836–42.

55. Fu L, Weiskopf EN, Akkermans O, Swanson NA, Cheng S, Schwartz TU, et al. HIV-1 capsids enter the FG phase of nuclear pores like a transport receptor. Nature. 2024;626(8000):843–51.

56. Price AJ, Jacques DA, McEwan WA, Fletcher AJ, Essig S, Chin JW, et al. Host Cofactors and Pharmacologic Ligands Share an Essential Interface in HIV-1 Capsid That Is Lost upon Disassembly. PLOS Pathogens. 2014;10(10):e1004459.

57. Lamorte L, Titolo S, Lemke Christopher T, Goudreau N, Mercier J-F, Wardrop E, et al. Discovery of Novel Small-Molecule HIV-1 Replication Inhibitors That Stabilize Capsid Complexes. Antimicrobial Agents and Chemotherapy. 2013;57(10):4622–31.

58. Bester SM, Wei G, Zhao H, Adu-Ampratwum D, Iqbal N, Courouble VV, et al. Structural and mechanistic bases for a potent HIV-1 capsid inhibitor. Science. 2020;370(6514):360–4.

59. Bruce A, Adebomi V, Czabala P, Palmer J, McFadden WM, Lorson ZC, et al. A Tag-Free Platform for Synthesis and Screening of Cyclic Peptide Libraries. Angewandte Chemie International Edition. 2024;63(21):e202320045.

60. Kirby KA, McFadden WM, Wang L, Du H, Zhang H, Emanuelli Castaner A, et al. Structural, biophysical, and virological mechanistic characterization of HIV-1 capsid-targeting antivirals. bioRxiv. 2025:2025.12.30.696960.

61. Jang S, Engelman Alan N. Capsid–host interactions for HIV-1 ingress. Microbiology and Molecular Biology Reviews. 2023;87(4):e00048–22.

62. Zhuang S, Torbett BE. Interactions of HIV-1 Capsid with Host Factors and Their Implications for Developing Novel Therapeutics. Viruses. 2021;13(3):417.

63. Wang L, Casey MC, Vernekar SKV, Do HT, Sahani RL, Kirby KA, et al. Chemical profiling of HIV-1 capsid-targeting antiviral PF74. European Journal of Medicinal Chemistry. 2020;200:112427.

64. Pennetzdorfer N, Naik V, Demirdjian S, Hendricks MR, Jamieson CS, Perry JK, et al. Lenacapavir treatment–emergent HIV-1 capsid resistance mutations are frequently associated with replication defects. Science Translational Medicine. 2026;18(831):eaea0947.

65. Jacques DA, McEwan WA, Hilditch L, Price AJ, Towers GJ, James LC. HIV-1 uses dynamic capsid pores to import nucleotides and fuel encapsidated DNA synthesis. Nature. 2016;536(7616):349–53.

66. Dick RA, Zadrozny KK, Xu C, Schur FKM, Lyddon TD, Ricana CL, et al. Inositol phosphates are assembly co-factors for HIV-1. Nature. 2018;560(7719):509–12.

67. Mallery DL, Márquez CL, McEwan WA, Dickson CF, Jacques DA, Anandapadamanaban M, et al. IP6 is an HIV pocket factor that prevents capsid collapse and promotes DNA synthesis. eLife. 2018;7:e35335.

68. Piacentini J, Allen DS, Ganser-Pornillos BK, Chanda SK, Yoh SM, Pornillos O. Molecular Determinants of PQBP1 Binding to the HIV-1 Capsid Lattice. J Mol Biol. 2024;436(4):168409.

69. Yoh SM, Mamede JI, Lau D, Ahn N, Sánchez-Aparicio MT, Temple J, et al. Recognition of HIV-1 capsid by PQBP1 licenses an innate immune sensing of nascent HIV-1 DNA. Molecular Cell. 2022;82(15):2871–84.e6.

70. Renner N, Mallery DL, Faysal KMR, Peng W, Jacques DA, Böcking T, et al. A lysine ring in HIV capsid pores coordinates IP6 to drive mature capsid assembly. PLOS Pathogens. 2021;17(2):e1009164.

71. McFadden WM, Gao X, Ye Z, Wen X, Lorson ZC, Zheng H, et al. TSAR, “Thermal Shift Analysis in R”, identifies endogenous molecules that interact with HIV-1 capsid hexamers. bioRxiv. 2025:2023.11.29.569293.

72. Kleinpeter A, Mallery DL, Renner N, Albecka A, Klarhof JO, Freed EO, et al. HIV-1 adapts to lost IP6 coordination through second-site mutations that restore conical capsid assembly. Nature Communications. 2024;15(1):8017.

73. Huang S-W, Briganti L, Annamalai AS, Greenwood J, Shkriabai N, Haney R, et al. The primary mechanism for highly potent inhibition of HIV-1 maturation by lenacapavir. PLOS Pathogens. 2025;21(1):e1012862.

74. dos Santos NFB, Lewis JA, Hansen M, Pereira MJB, Christensen DE, Sundquist WI, et al. Lenacapavir allosterically remodels the HIV-1 capsid. bioRxiv. 2026:2026.01.05.697065.

75. Gupta M, Waltmann C, Renner N, Wang Y, James LC, Jacques DA, et al. Mechanistic insights into lenacapavir-induced off-pathway HIV-1 capsid assembly. Proceedings of the National Academy of Sciences. 2026;123(11):e2524995123.

76. McFadden WM, Kirby KA, Wang L, Du H, Lorson ZC, Zhang H, et al. Modifying PF74 Improves Anti-HIV-1 Activity Against the Resistance-Associated Capsid Mutation N74D. [CROI Abstract 727]. Topics in Antiviral Medicine. 2025;33(1):208.

77. Zhou J, Price Amanda J, Halambage Upul D, James Leo C, Aiken C. HIV-1 Resistance to the Capsid-Targeting Inhibitor PF74 Results in Altered Dependence on Host Factors Required for Virus Nuclear Entry. Journal of Virology. 2015;89(17):9068–79.

78. Saito A, Ferhadian D, Sowd Gregory A, Serrao E, Shi J, Halambage Upul D, et al. Roles of Capsid-Interacting Host Factors in Multimodal Inhibition of HIV-1 by PF74. Journal of Virology. 2016;90(12):5808–23.

79. Rankovic S, Ramalho R, Aiken C, Rousso I. PF74 Reinforces the HIV-1 Capsid To Impair Reverse Transcription-Induced Uncoating. Journal of Virology. 2018;92(20):e00845–18.

80. Lad L, Clancy S, Koditek D, Wong MH, Jin D, Niedziela-Majka A, et al. Functional Label-Free Assays for Characterizing the in Vitro Mechanism of Action of Small Molecule Modulators of Capsid Assembly. Biochemistry. 2015;54(13):2240–8.

81. Lanman J, Sexton J, Sakalian M, Prevelige PE, Jr. Kinetic analysis of the role of intersubunit interactions in human immunodeficiency virus type 1 capsid protein assembly in vitro. J Virol. 2002;76(14):6900–8.

82. McFadden WM, Casey-Moore MC, Bare GAL, Kirby KA, Wen X, Li G, et al. Identification of clickable HIV-1 capsid-targeting probes for viral replication inhibition. Cell Chemical Biology. 2024;31(3):477–86.e7.

83. Sticht J, Humbert M, Findlow S, Bodem J, Müller B, Dietrich U, et al. A peptide inhibitor of HIV-1 assembly in vitro. Nature Structural & Molecular Biology. 2005;12(8):671–7.

84. Miyazaki Y, Doi N, Koma T, Adachi A, Nomaguchi M. Novel In Vitro Screening System Based on Differential Scanning Fluorimetry to Search for Small Molecules against the Disassembly or Assembly of HIV-1 Capsid Protein. Front Microbiol. 2017;8(1413):1413.

85. Schirra RT, dos Santos NFB, Ganser-Pornillos BK, Pornillos O. Arg18 Substitutions Reveal the Capacity of the HIV-1 Capsid Protein for Non-Fullerene Assembly. Viruses. 2024;16(7):1038.

86. Briganti L, Annamalai Arun S, Bester Stephanie M, Wei G, Andino-Moncada Jonathan R, Singh Satya P, et al. Structural and mechanistic bases for resistance of the M66I capsid variant to lenacapavir. mBio. 2025;16(5):e03613–24.

87. Pornillos O, Ganser-Pornillos BK, Yeager M. Atomic-level modelling of the HIV capsid. Nature. 2011;469(7330):424–7.

88. Zhao G, Perilla JR, Yufenyuy EL, Meng X, Chen B, Ning J, et al. Mature HIV-1 capsid structure by cryo-electron microscopy and all-atom molecular dynamics. Nature. 2013;497(7451):643–6.

89. Pettersen EF, Goddard TD, Huang CC, Meng EC, Couch GS, Croll TI, et al. UCSF ChimeraX: Structure visualization for researchers, educators, and developers. Protein Science. 2021;30(1):70–82.

90. Fu L, Cheng S, Riedel D, Kopecny L, Schuh M, Görlich D. Nuclear pore passage of the HIV capsid is driven by its unusual surface amino acid composition. Nature Structural & Molecular Biology. 2025;32(12):2476–91.

91. Kimanius D, Dong L, Sharov G, Nakane T, Scheres SHW. New tools for automated cryo-EM single-particle analysis in RELION-4.0. Biochemical Journal. 2021;478(24):4169–85.

92. Punjani A, Rubinstein JL, Fleet DJ, Brubaker MA. cryoSPARC: algorithms for rapid unsupervised cryo-EM structure determination. Nature Methods. 2017;14(3):290–6.

93. Sterling HJ, Prell JS, Cassou CA, Williams ER. Protein conformation and supercharging with DMSO from aqueous solution. J Am Soc Mass Spectrom. 2011;22(7):1178–86.

94. Harvey S, editor Characterizing the interactions between the HIV-1 capsid protein and small molecule ligands using native mass spectrometry. PROTEIN SCIENCE; 2024: WILEY 111 RIVER ST, HOBOKEN 07030-5774, NJ USA.

95. Harvey SR, O’Neale C, Schey KL, Wysocki VH. Native Mass Spectrometry and Surface Induced Dissociation Provide Insight into the Post-Translational Modifications of Tetrameric AQP0 Isolated from Bovine Eye Lens. Analytical Chemistry. 2022;94(3):1515–9.

