## Supplementary Table S1 and Figures for "Damaging the conical morphology of HIV-1 capsid by targeting the FG-binding pocket and disfavoring pentameric subunits needed for core closure"

| | | **T1:1261** | **T1 Apo** | **CLP:1261[a]** |
|---|---|---|---|---|
| **Sample details** | Sample pH | 8.0 | 8.0 | 8.0 |
| | IP6 | + | + | + |
| **Data collection** | Microscope | Talos Arctica | Talos Arctica | Talos Arctica |
| | Detector | K3 (Gatan) | K3 (Gatan) | K3 (Gatan) |
| | Nominal magnification | 79,000x | 79,000x | 63,000x |
| | Voltage (kV) | 200 | 200 | 200 |
| | Total dose ($e^{-}$/Å$^{2}$) | 48.7 | 50.8 | 27.7 |
| | Super-resolution mode? | yes | yes | yes |
| | Acquisition software | EPU | EPU | EPU |
| | Defocus range (µm) | -0.6 to -1.8 | -0.6 to -1.8 | ~ -0.5 to -2.0 |
| | Pixel size (Å/px) | 1.08 | 1.08 | 1.31 |
| | Frames per movie | 50 | 50 | 50 |
| | Number of movies | 2,584 | 2,022 | 3,437 |
| **Processing** | Final number of particles | 621,749 | 83,621 | 120,885 |
| | Symmetry imposed | I | I | C6 |
| | Map resolution at 0.143 FSC (Å) | 2.32 | 2.74 | 4.76 |
| **Atomic model** | Protein residues | 221 | 221 | - |
| | MolProbity score | 1.81 | 1.77 | - |
| | Clash score | 7.54 | 8.18 | - |
| | Rotamer outliers (%) | 1.59 | 2.65 | - |
| | Ramachandran favored (%) | 96.35 | 98.63 | - |
| | Ramachandran allowed (%) | 3.65 | 1.37 | - |
| | Ramachandran outliers (%) | 0.00 | 0.00 | - |
| | RMSD, bond length (Å) | 0.004 (0) | 0.003 (0) | - |

| | | | | |
|---|---|---|---|---|
| | RMSD, bond angles (°) | 0.909 (1) | 0.620 (0) | - |

**Supplementary Table S1: Summary of Cryo-EM Experiments**

[a]CLP:1261 data did not include protein modeling.

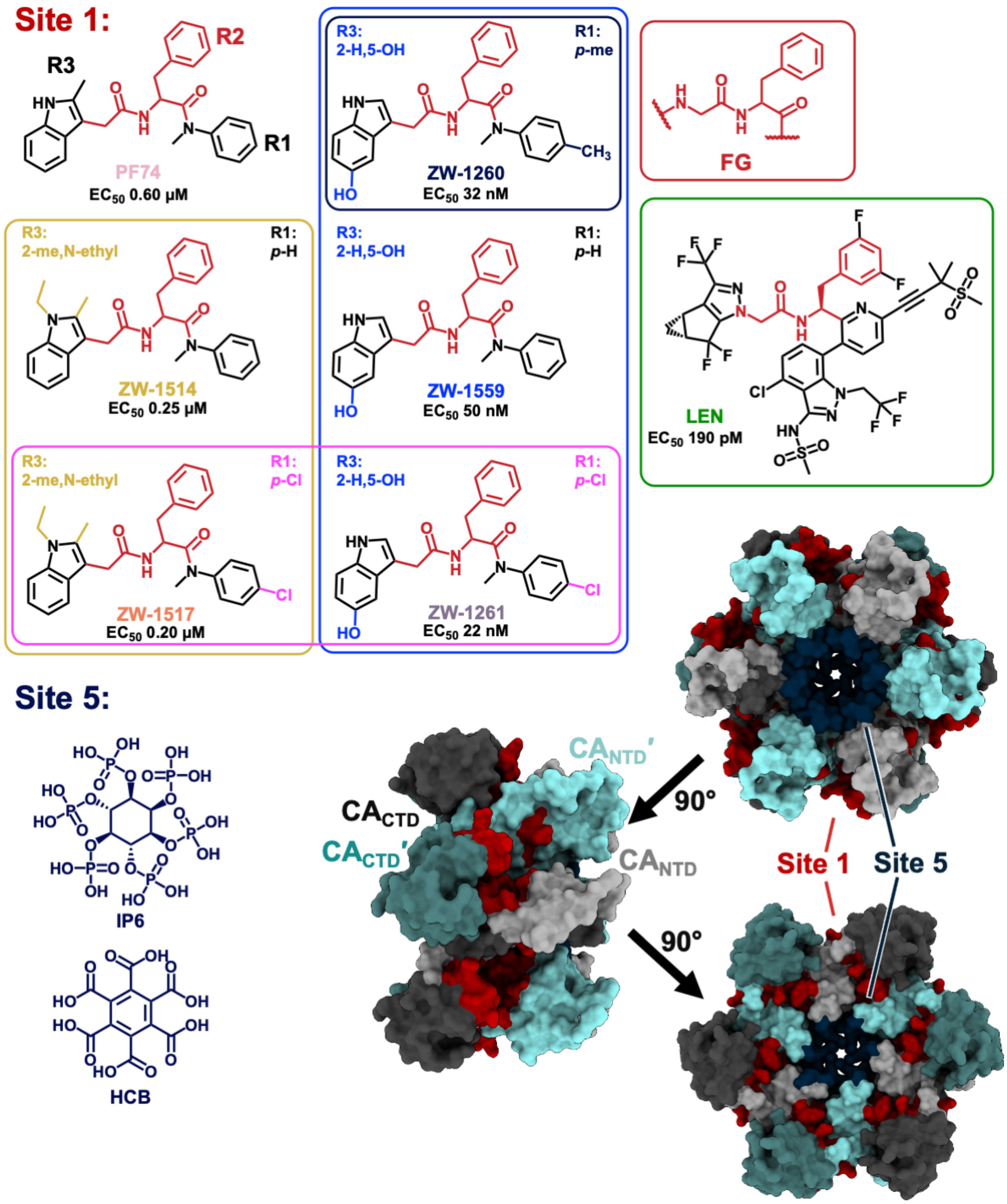


**Supplementary Figure S1: Phenylalanine-Glycine (FG) analog-containing antivirals and other compounds used in this study bind to CA Site 1 or Site 5.**

PF74 (Pink) was the first compound reported to bind at this site (16) and has three rings (R1, R2, and R3). ZW-1514 (Yellow) and ZW-1517 (Orange) modify the R3 ring with an ethyl group adjacent the methyl. ZW-1559 (Blue), ZW-1260 (Navy Blue), and ZW-1261 (Purple) modify the R3 indole to remove the methyl group on one side and add an alcohol group to the other ring in the indole. ZW-1260, ZW-1261, and ZW-1517 have an additional modification on the R1 ring, with ZW-1260 having a 4-methyl group while ZW-1261 and ZW-1517 have a 4-Cl. Lenacapavir (LEN, Green) is the only approved drug that targets the same site (4). Structures and $EC_{50}$ values from (10, 14, 16, 29, 63). These compounds and others with the FG-scaffold may bind to the FG-binding pocket shared by neighboring protomers in HIV-1 CA hexamers, shaded in red; the central pore for IP6 binding is in navy [PDB: 4XFX (42)]. CA monomers are depicted with the N-terminal domain (NTD) in lighter shades and the C-terminal domain (CTD) in darker shades. IP6 and hexacarboxybenzene (HCB) are shown below that bind site 5, the central pore.

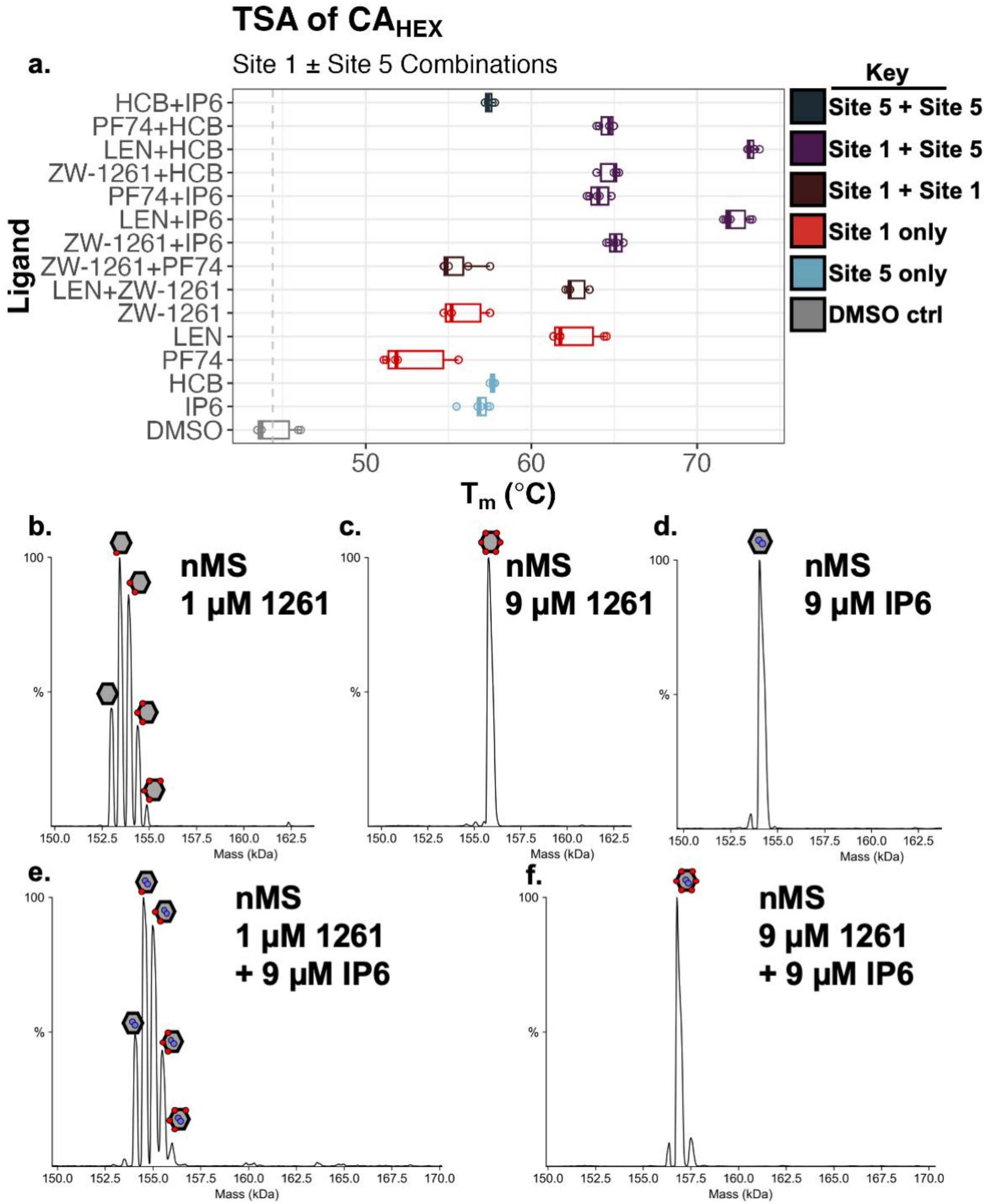


**Supplementary Figure S2: FG-binding pocket can be saturated with one compound, and is further stabilized by IP6.**

**A.** TSA of $CA_{HEX}$ treated with FG-binding compounds ± Pore-binding compounds. The two different sites have additive effect on the $\Delta T_m$ for $CA_{HEX}$. Boxplot produced with TSAR (71). **B-F.** Native Mass Spectrometry (nMS) analysis of $CA_{HEX}$ finds ZW-1261 **B.** can bind multiple pockets at 1 μM and **C.** fully saturate a $CA_{HEX}$ by binding to all six pockets at 9 μM. **D.** For a $CA_{HEX}$, two IP6 molecules bind to the central pore at 9 μM IP6. **E**. A combination of 1 μM ZW-1261 and 9 μM IP6 find two IP6 bound and between one and four ZW-1261 bound to a $CA_{HEX}$. **F.** Both IP6 and ZW-1261 can saturate a $CA_{HEX}$ at 9 μM IP6 + 9 μM ZW-1261.

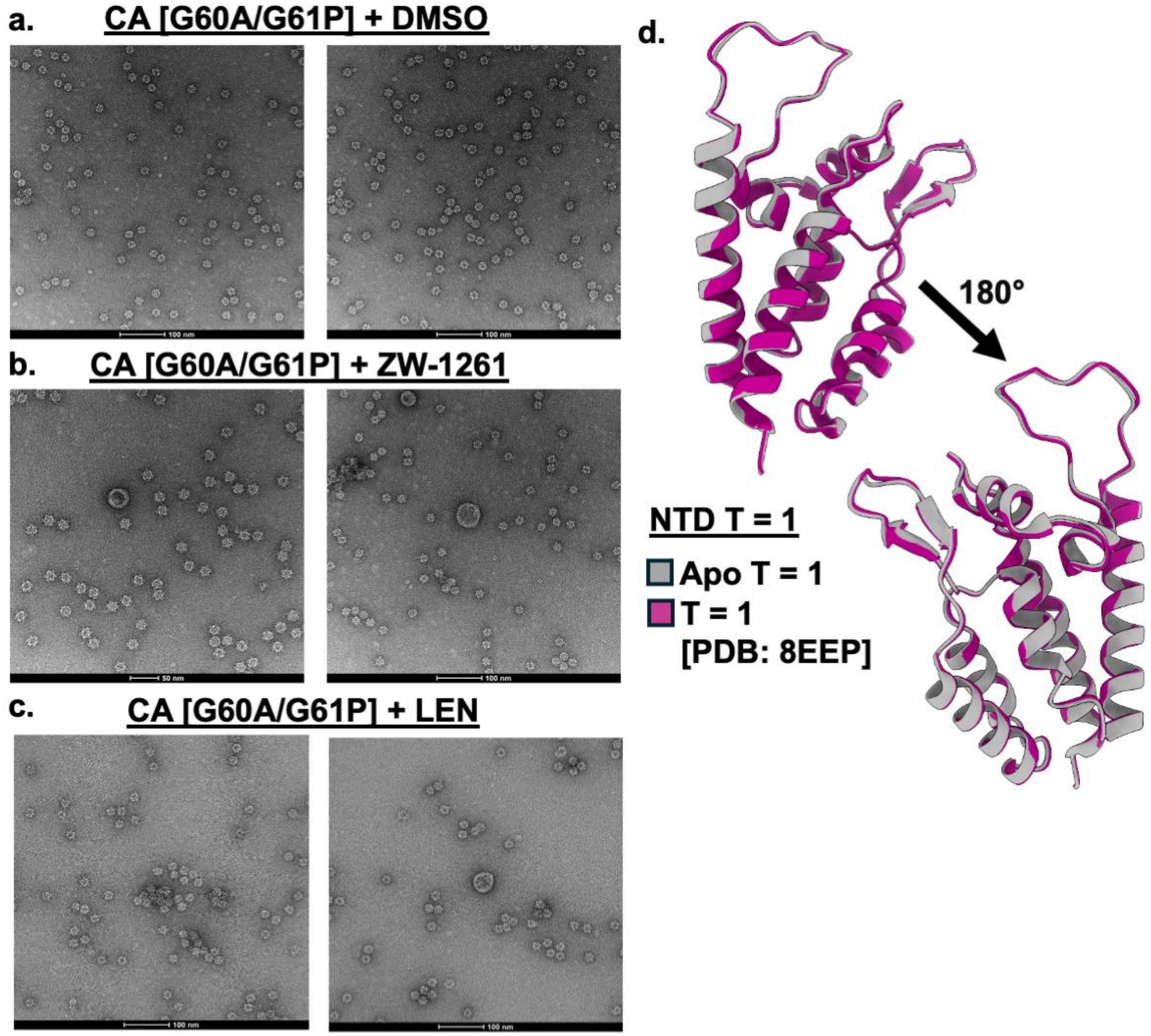


**Supplementary Figure S3: Negative stain TEM images of T = 1 particles treated with FGBP-targeting antivirals.**

**A.** A pentameric-only assembly of CA forms with mutations G60A/G61P that change the TVGG switch and make a particle with 12 pentamers and 20 nm diameter (41). **B.** Treating pre-formed T = 1 CA particles from (**A**) with ZW-1261 leads to increased aggregation as well as occasionally larger or damaged assemblies. **C.** Same as (**B**), treating T = 1 CA with LEN leads to leads to apparent differences in morphology. **D.** We solved the T = 1 Icosahedral $CA_{PENT}$-only construct in the absence of inhibitor (grey) at 2.74 Å and find the TVGG loop in the untreated condition is identical to the previously reported structure (pink) [PDB: 8EEP (41)]. Thus, adding a HIS-tag to the $CA_{CTD}$ does not impact the FGBP.
